# An in vivo and in silico evaluation of the magical hepatoprotective potentialities of Gynura procumbens: a promising agent for combating hepatotoxicity

**DOI:** 10.1101/2022.05.11.491443

**Authors:** Tanzia Islam Tithi, Md. Rafat Tahsin, Tasnuva Sharmin Zaman, Juhaer Anjum, Nasiba Binte Bahar, Priyanka Sen, Sabiha Tasnim, Arifa Sultana, Fahima Jannat Koly, Ishrat Jahan, Fahima Aktar, Jakir Ahmed Chowdhury, Shaila Kabir, Abu Asad Chowdhury, Md. Shah Amran

**Affiliations:** Department of Pharmaceutical Technology, Faculty of Pharmacy, University of Dhaka, Dhaka 1000, Bangladesh; Department of Pharmaceutical Sciences, North South University, Plot # 15, Block # B, Bashundhara R/A, Dhaka 1229, Bangladesh; Department of Pharmacy, Faculty of Pharmacy, University of Dhaka, Dhaka 1000, Bangladesh; Deartment of Clinical Pharmacy and Pharmacology, Faculty of Pharmacy, University of Dhaka, Dhaka 1000, Bangladesh; Molecular Pharmacology and Herbal Drug Research Laboratory, Department of Pharmaceutical Chemistry, Faculty of Pharmacy, University of Dhaka, Dhaka 1000; Department of Pharmacy, University of Asia Pacific, Farmgate, Dhaka, Bangladesh, University of Dhaka, Dhaka 1000

**Keywords:** hepatotoxicity, *Gynura procumbens*, hepatoprotective, liver

## Abstract

**Introduction:** Liver being the most important metabolic organ of the body performs a wide variety of vital functions. Hepatic cell injury occurs by the activation of reactive oxygen species (ROS) by CCl_4_, xenobiotics and other toxic substances generated through cytochrome P450 dependent step resulting from covalent bond formation with lipoproteins and nucleic acids. Observing the alarming state of hepatotoxic patients worldwide, different medicinal plants and their properties can be explored to combat against such free radical degermation of liver. This paper evaluates the antioxidant property of *Gynura procumbens* in both in silico and in an in vivo assay, and its hepatoprotective activity in CCl_4_ induced hepatotoxicity.

**Materials and Methods:** *Gynura procumbens* leaves were collected and extracted using 50% ethanol. Required chemicals (CCl_4_), standard drug (Silymarin) and blood serum analyzing kits were stocked. The in vivo tests were performed in 140 healthy Wister albino male rats under well controlled parameters dividing into 14 groups, strictly maintaining IEAC protocols. In silico molecular docking and ADMET studies were performed and the results were analyzed statistically.

**Results and discussion:** The body weight increased significantly in CCl_4_ induced, *G. procumbens* administered hepatotoxic rats. The increase in SGPT, SGOT, ALP, creatinine, LFH, triglycerides, LDL, SOD, MDA, total cholesterol, DNA fragmentation ranges, γGT levels of CCl_4_ treated group was decreased by both standard drug Silymarin and *G. procumbens* leaf extract. On the other hand, *G. procumbens* increased HDL levels and displayed contrasting results in CAT level tests. Some results contradicted with the negative controlled group displaying varying efficacy between leaf extract and Silymarin. In the molecular docking analysis, *G. procumbens* phytoconstituents performed poorly against TGF-β1 compared to the control drug Galunisertib while 26 phytoconstituents scored better than the control, bezafibrate against PPAR-α. Flavonoids and phenolic compounds performed better than other constituents in providing hepatoprotective activity.

## Introduction

Liver is a very important and resilient organ in an animal body, major function of which is to give support in metabolism, digestion, detoxification, storing vitamins, and immunity. However, this organ can be affected by different toxins, which originates commonly from food supplements, drugs, chemicals, and medicinal plants [1–3]. Carbon tetra chloride (CCl_4_) is considered as quite toxic and as well as xenobiotic to animal body that causes the cell injury by activation of reactive oxygen species (ROS). Peroxy trichloromethyl (OOCCl_3_) and trichloromethyl (CCl_3_) radicals are usually generated during cytochrome P450 dependent step. These free radicals produce covalent bond with lipoproteins and nucleic acids that cause extensive cell injury in liver and other important organs of the animal body [4–10]. Till date, around fifty million people all throughout the world are suffering from hepatotoxicity. This state of hepatotoxicity is quite alarming in health sector [1]. Symptoms that are related to hepatotoxicity may encompass jaundice, fatigue, abdominal pain, skin rashes, itching, rapid and abnormal weight gain, vomiting or nausea, light colored stool and dark urine [11, 12]. To examine liver function among different diagnostic systems, serum glutamate pyruvate transaminase (SGPT), serum glutamate oxaloacetate transaminase (SGOT), acid phosphatase (ACP), and alkaline phosphatase (ALP) level are measured in the blood [13, 14]. Different synthetic drugs such as, thalidomide, curcumin, ademethionine, entecavir, metadoxine, tenofovir, ondensatron, and resveratrol are available in the pharma arena, which are used to treat liver diseases [15]. However, the toxicity and efficacy of these medicines are still under investigation, where medicinal plants have great opportunity to take their position by genetic modification using different biotechnological studies [16]. Medicinal plants have always had a great impact on medicines as well as nutrition for different societies in the world. It is estimated that, around 80,000 medicinal plants are available in this planet [17]. Not only in developing countries like China, India, Pakistan and Sri Lanka (more than 80%) but also in developed country such as United States (more than 25%) drugs are constituted from medicinal plants [18–19]. More specifically, *Gynura procumbens* is a very special herb which is widely available in south-east region of the world such as; Thailand, Malaysia, China, Vietnam, Bangladesh and Indonesia. [20]. It is a small (around 1-3 m long) flowering plant which belongs to Asteraceae family. This herb is taken as a vegetable item in meals and also used as medicinal plant in these regions. The major pharmacological activities of *G. Procumbens* are on diabetes, Herpes Simplex Virus (HSV), hyperlipidemia, inflammation, colon cancer, hemorrhoids, analgesic, kidney disease, rheumatism and hypertension [21–24]. In addition, different bioactive molecules such as, saponins, sterol, tannins, terpenoids, flavonoids, and glycoside are useful chemical components to treat these sorts of diseases successfully. Moreover, the most recent revelation of antioxidant properties of this plant gives a clear and precise idea to treat various diseases by the reduction of free radical production [25–28]. For this reason, the local name of this plant is “Pokok Sambung Nyawa” the meaning of which is “prolongation of life” [29–30]. The main aim of this manuscript is to assay the hepatoprotective activities of *G. procumbens* in vivo in carbon tetra cloride (CCl_4_)-induced liver toxicity and in silico and to computationally predict the ADMET properties of the phytoconstituents.

## Method and Materials

### Collection and extraction of *Gynura procumbens* leaf

First of all, the *Gynura procumbens* leaf was collected from the garden of the Pharmacy department, Jahangir Nagar University. After that, the specimen was certified by the Bangladesh National herbarium and provided the accession number YM001 for the future references. The wet *Gynura procumbens* leaf was carefully powdered coarsely after air drying. After that, the extraction of powdered leaf was done by using 50% ethanol for few days. The extract was filtered after every 3 days. Later on, rotary evaporator was used to dry the extract using low temperature and pressure. Finally, different pharmacological tests were performed using this crude residue.

### Drugs and Chemicals

Carbon tetra chloride (CCl_4_) was bought from the Sigma Aldrich company, USA. Standard hepatoprotective drug named Silymarin (Brand Name: Silybin) was purchased from Square Pharmaceuticals Ltd. Again, the blood serum analyzing kits, SGPT, SGOT, ALP, γ-GT, Creatinine, Urea, Total Cholesterol, HDL, LDL, Triglyceride, CAT, SOD and MDA were bought from Plasmatec Laboratory Product Ltd. Bangladesh.

### Animals

140 healthy male rats (Wistar rats) weighing between (150-200) gm was purchased from North South University, Dhaka and each of them was nourished carefully by keeping them in the Institute of Nutrition & Food Science at the University of Dhaka in a well-controlled environment (relative humidity 55±5%, 12±1h light/dark cycle and temperature 25±3°C) for 2 weeks. After that, considering equal body mass index, 14 groups were created where each group was constituted with 10 rodents. All rats were provided with a standard food supplement and filtered water. Finally, all of the experimental procedures were carried out according to the Institutional Animals Ethics Committee (IEAC).

**Table 1:**
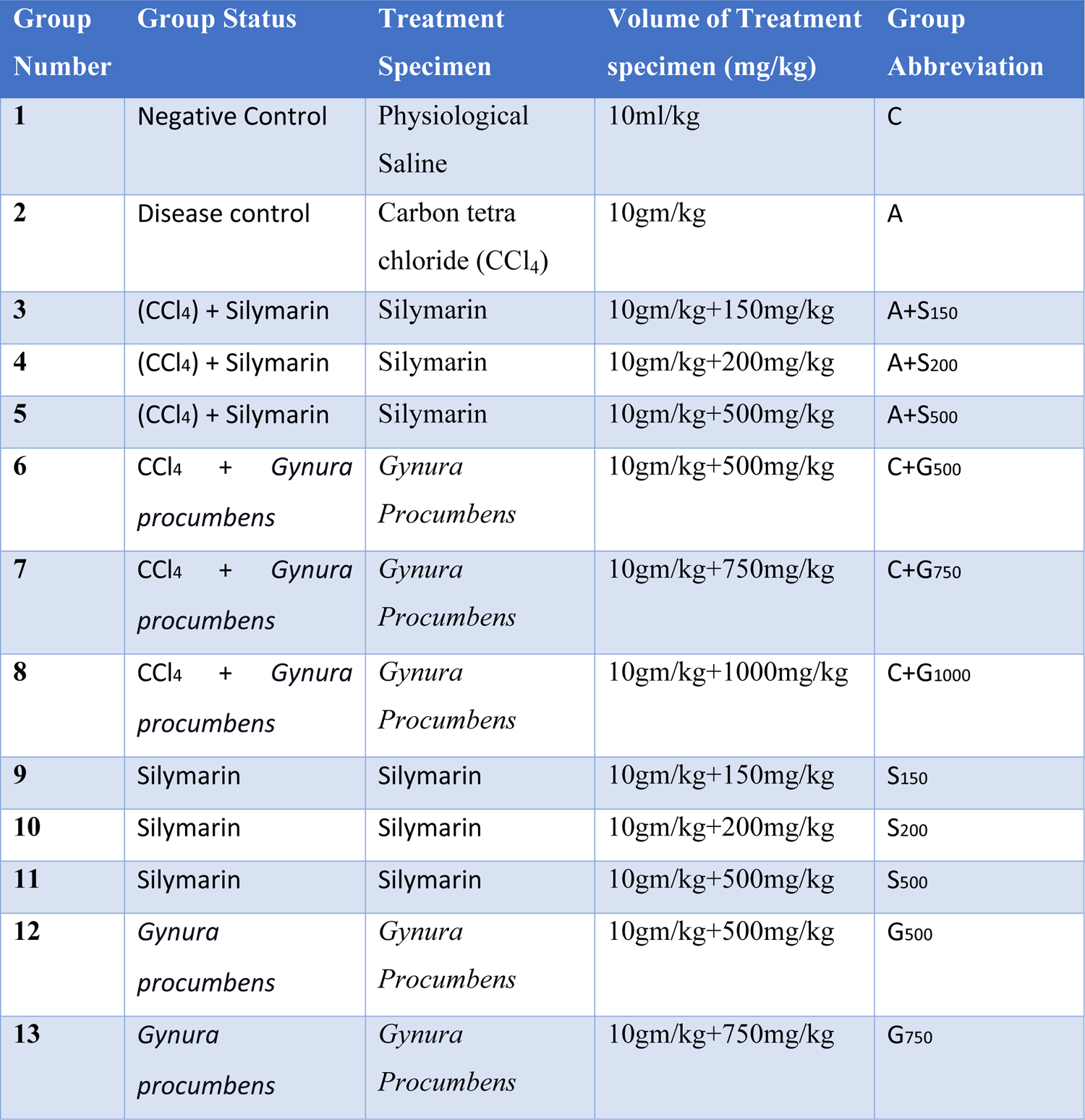
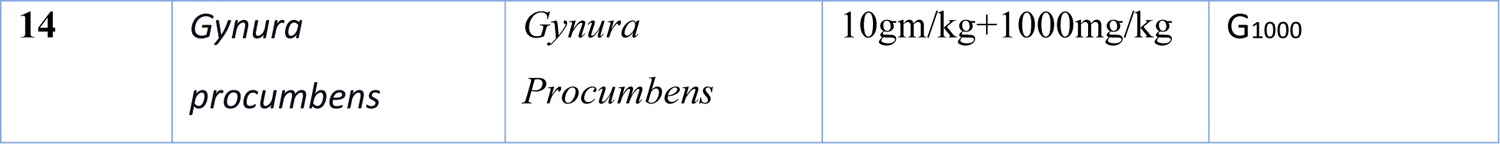
Evaluation of Hepatoprotective Activity

| Group Number | Group Status | Treatment Specimen | Volume of Treatment specimen (mg/kg) | Group Abbreviation |
| --- | --- | --- | --- | --- |
| 1 | Negative Control | Physiological Saline | 10ml/kg | C |
| 2 | Disease control | Carbon tetra chloride (CCl<sub>4</sub>) | 10gm/kg | A |
| 3 | (CCl<sub>4</sub>) + Silymarin | Silymarin | 10gm/kg+150mg/kg | A+S<sub>150</sub> |
| 4 | (CCl<sub>4</sub>) + Silymarin | Silymarin | 10gm/kg+200mg/kg | A+S<sub>200</sub> |
| 5 | (CCl<sub>4</sub>) + Silymarin | Silymarin | 10gm/kg+500mg/kg | A+S<sub>500</sub> |
| 6 | CCl<sub>4</sub> + *Gynura procumbens* | *Gynura Procumbens* | 10gm/kg+500mg/kg | C+G<sub>500</sub> |
| 7 | CCl<sub>4</sub> + *Gynura procumbens* | *Gynura Procumbens* | 10gm/kg+750mg/kg | C+G<sub>750</sub> |
| 8 | CCl<sub>4</sub> + *Gynura procumbens* | *Gynura Procumbens* | 10gm/kg+1000mg/kg | C+G<sub>1000</sub> |
| 9 | Silymarin | Silymarin | 10gm/kg+150mg/kg | S<sub>150</sub> |
| 10 | Silymarin | Silymarin | 10gm/kg+200mg/kg | S<sub>200</sub> |
| 11 | Silymarin | Silymarin | 10gm/kg+500mg/kg | S<sub>500</sub> |
| 12 | *Gynura procumbens* | *Gynura Procumbens* | 10gm/kg+500mg/kg | G<sub>500</sub> |
| 13 | *Gynura procumbens* | *Gynura Procumbens* | 10gm/kg+750mg/kg | G<sub>750</sub> |
| 14 | *Gynura procumbens* | *Gynura Procumbens* | 10gm/kg+1000mg/kg | G<sub>1000</sub> |

Group 2-8 was administered with carbon tetra chloride (CCl_4_) at a dose of 10 mg/kg through an intraperitoneal route to induce hepatotoxicity. Then different parameters such as; SGPT, SGOT, ALP, γ-GT, Creatinine, Urea, Total Cholesterol, HDL, LDL, Triglyceride, CAT, SOD and MDA were measured in all groups to check whether liver cirrhosis was induced or not. The duration of treatment was six weeks. Both the drugs and the extracts were given orally.

### Statistical Analysis

All our findings (raw data) belong to several groups regarding numerous research parameters recorded and analyzed on a broadsheet using MS excel program. Data were subjected to descriptive statistics and results were represented as mean ± SD. We employed the “One Way Anova Test’’ of SPSS 16” software for interpreting the inter-group heterogeneity in terms of diverse biological parameters to determine the statistical significance. We consider the events as statistically significant while the ‘p’ value was detected as less than 0.05 (p<0.5).

### In silico molecular docking and ADMET studies

76 *G. procumbens* phytoconstituents obtained from literature mining comprised of the ligand library for the in-silico assessment of hepatoprotectivity. 3D co-ordinate files of the constituents were either downloaded from the ‘PubChem’ or the ‘ChemSpider’ databases or drawn manually on the ‘Avogadro’ software package [31, 32]. The macromolecules TGF-β Type I Receptor (PDB ID: 1VJY) and PPAR-α (PDB ID: 5HYK) were selected as the molecular targets for the docking studies [33–36]. The active sites of the macromolecules were deduced from literature, and galunisertib and bezafibrate were employed as control molecules, respectively [37–40]. The phytoconstituents and the control molecules were optimized under the MMFF94 forcefield employing the steepest descent algorithm (convergence value: 10e^-7^) using the software package ‘Avogadro’ and saved in the pdb format [41]. The macromolecules were downloaded from the ‘Protein Data Bank’ database and prepared using the software packages ‘PyMol’ and ‘Swiss-PdbViewer 4.1.0’, the latter of which was utilized to carry out energy minimization using the ‘GROMOS96’ forcefield with parameters set 43B1 in vacuo [42–45].

The AutoDock Vina component of the software package ‘PyRx’ was used to carry out molecular docking [46]. Results were visualized and analyzed in the software packages’ PyMol’ and ‘Discovery Studio Visualizer 2020’ [42, 47]. The canonical SMILES of the ligands were obtained using the software package ‘OpenBabel’ and used as inputs on the computational webservers ‘SwissADME’ and ‘ProTox-II’ to predict the ADMET properties of the phytoconstituents [48–50].

## Results

A significant elevation of body weight was observed after administering *Gynura procumbens* extract in CCl4 induced hapatotoxic rats [**Figure 1**]. In the negative control group, an escalation in body weight was obtained at the final stage. However, carbon tetrachloride treatment prompted a decrease in terminal body weight in the disease control group. Groups treating with 3 different doses of *Gynura procumbens* (Low, Medium, High) following carbon tetrachloride elevated the gradual pattern of weight-gain. However, silymarin treatment showed opposite scenario. A reversal of CCl_4_ mediated body weight decline was reported upon treatment with different doses of *Gynura procumbens* extract. Groups administered only plant extract followed a similar trend as the negative control group, whereas a contrasting scenario was observed in groups given Silymarin only. Increase in body weight after treating with following plant extracts are found in case of *Phytolacca dodecandra* [1].

**Figure 1.**
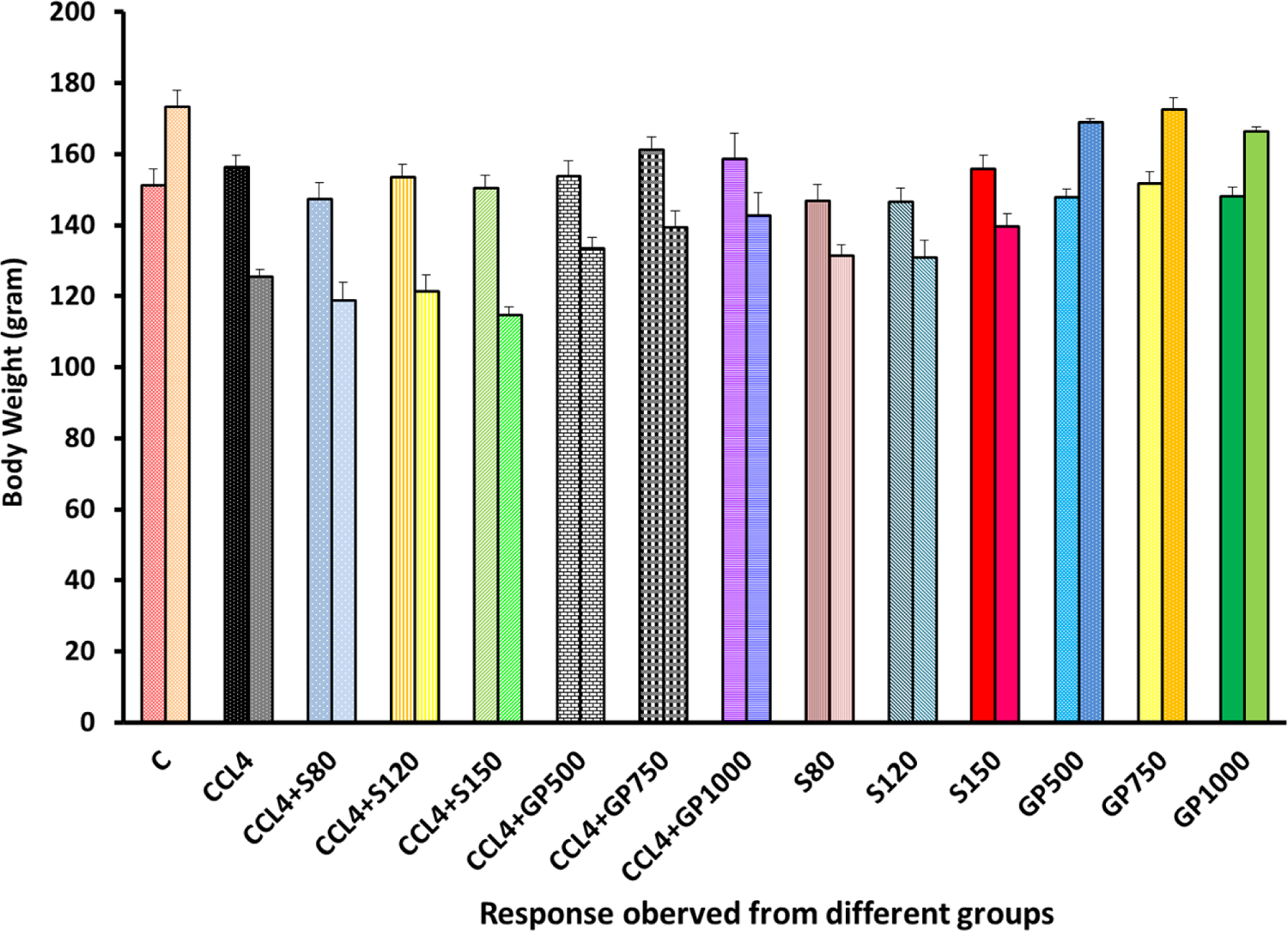
Body weight of rats of 14 groups before and after completing the experiment in liver affected rats.

The SGPT level of the carbon tetrachloride-treated group was increased to a higher level than that of the negative control. There was significant decline of SGPT level by both standard drug Silymarin and *Gynura procumbens* leaf extracts [**Figure 2**]. Similar types of decline in SGPT level was observed in some studies using leaf extracts of *Rhododendron arboreum*, *Beta vulgaris*, *Zanthoxylum armatum, Saururus chinensis, Curcuma longa, Limonium sinense, Hemidesmus indicus, Premna esculenta, Macrocybe gigantean*, and *Argyreia speciose* [2–11]. Although high doses of *Beta vulgaris* is needed to reduce the SGPT level in same amount as Silymarin [5].

**Figure 2:**
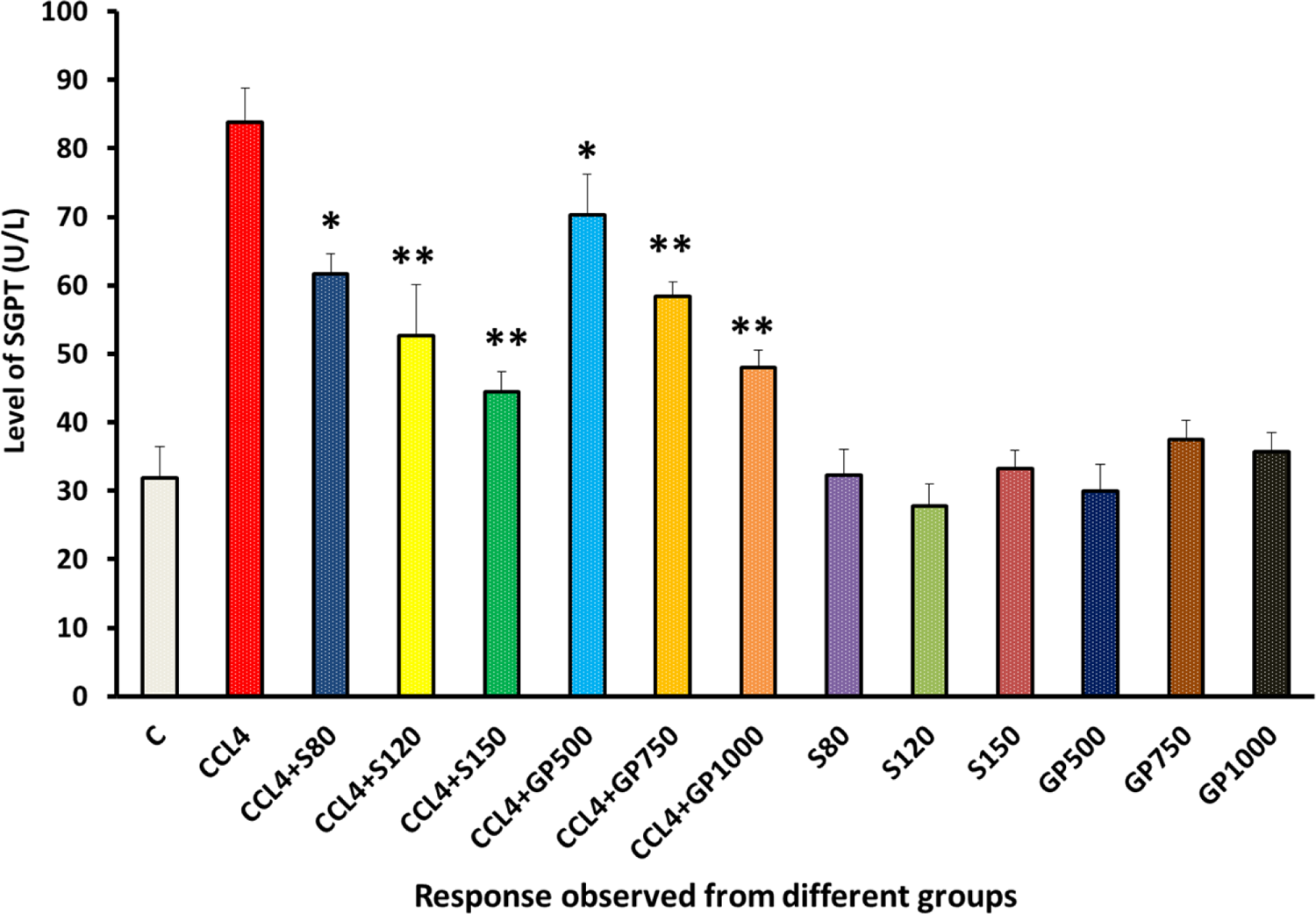
SGPT (U/L) Level of rats from 14 groups. The data were expressed as mean ± standard deviation. (* indicates statistically significant change)

SGOT level of positive control and disease control group showed two opposite scenarios [**Figure 3**]. Though SGOT levels were significantly reduced by both drugs and leaf extracts treated groups, Silymarin treated hepatotoxic rats showed better results than *Gynura procumbens* treated rats. Some studies with similar results have found using different plant extracts like-*Curcuma longa, Limonium sinense, Hemidesmus indicus, Rhododendron arboretum, Premna esculenta, Macrocybe gigantean*, and *Argyreia speciose* [2–4], [8–11].

**Figure 3:**
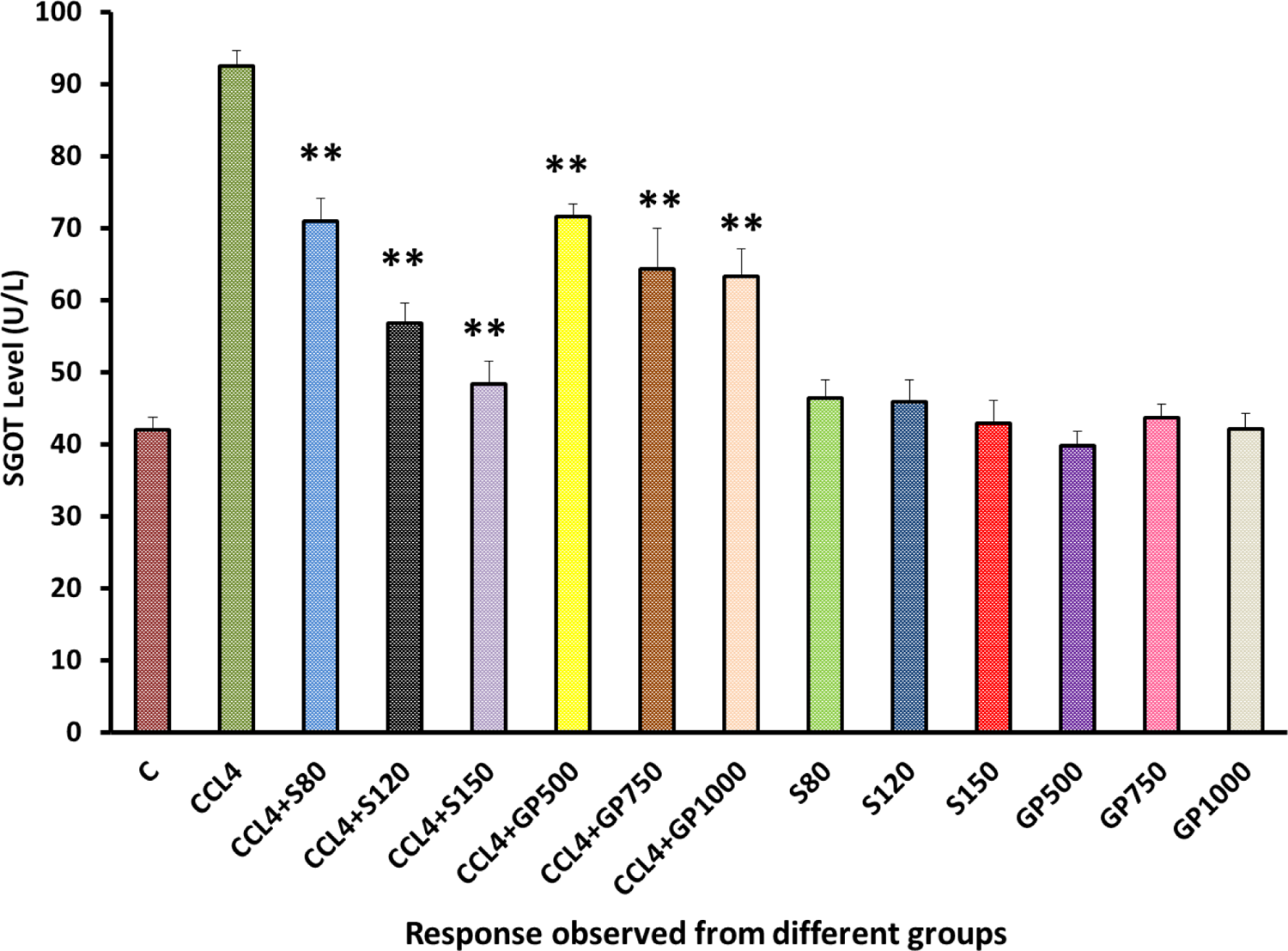
SGOT (U/L) Level of rats from 14 groups. The data were expressed as mean± standard deviation (* indicates statistically significant change)

Both drug and extracts showed statistical significance and ensured the effectiveness of *Gynura procumbens* extract [**Figure 4**]. No significant deviation could be spotted from the negative control group in the ALP level yielded by the remaining six non-carbon tetrachloride induced groups. Some studies have come to agreement with result for following plant extracts. *Pisonia aculeate, Solanum xanthocarpum, Beta vulgaris, Clerodendrum inerme, Rhododendron arboretum, Clutia abyssinica, Macrocybe gigantean, Argyreia speciose, Terminalia arjuna*, and *Phytolacca dodecandra* [1–5], [12–16].

**Figure 4:**
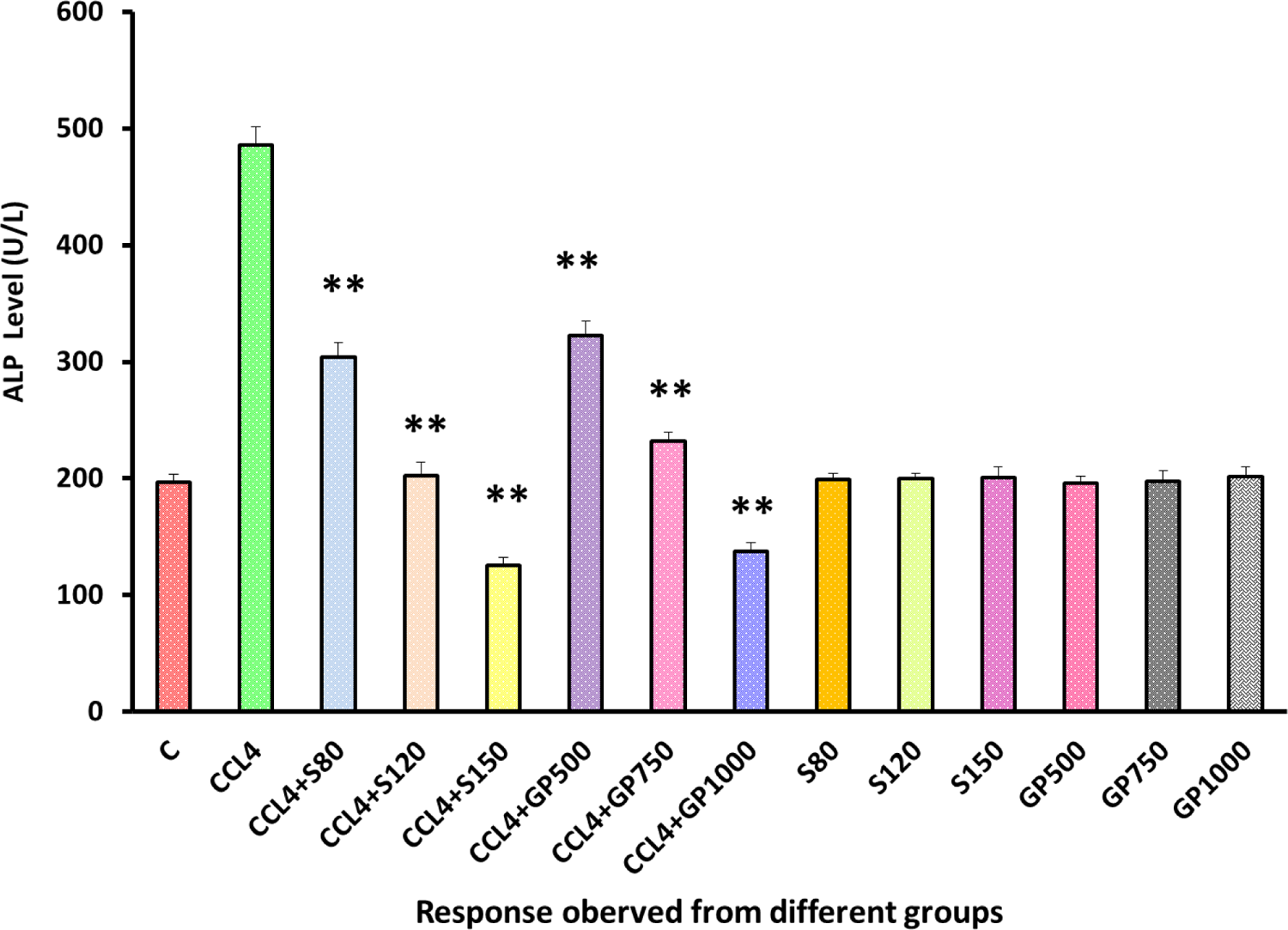
ALP level (U/L) Level of rats from 14 groups. The data were expressed as mean ± standard deviation. (* indicates statistically significant change)

A large increase in the creatinine level was displayed by the disease control group, which was in stark contrast with the negative control group. Silymarin and *Gynura procumbens* treatment gave prominent result by reducing creatinine level and there was no significant raise of this with negative control groups [**Figure 5**]. Creatinine level were noticed to be decreased by some other plant extracts like-*Syzygium aromaticum,* and *Marrubium vulgare* [17], [18].

**Figure 5:**
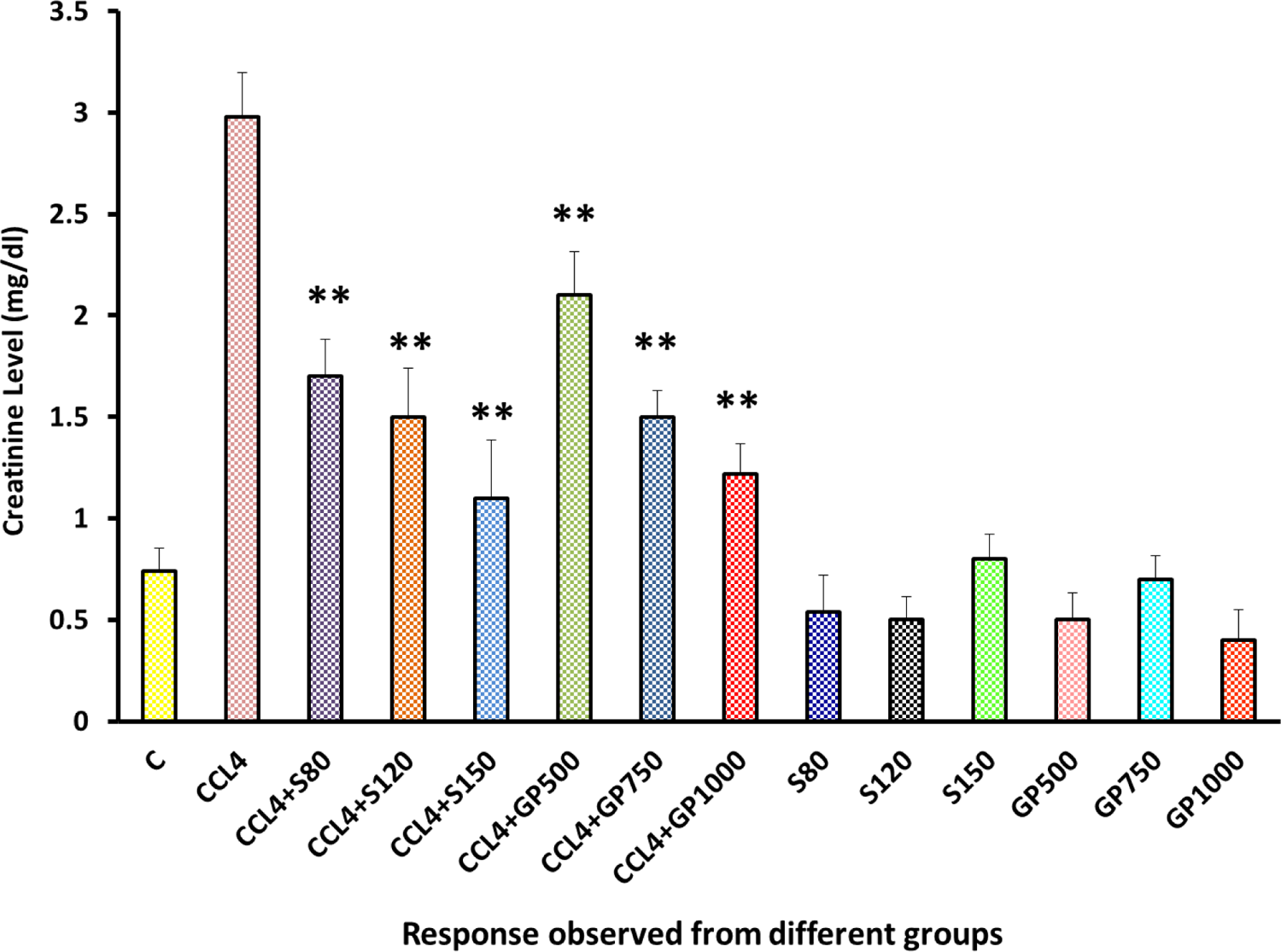
Creatinine (mg/dl) Level of rats from 14 groups. The data were expressed as mean ± standard deviation. (* indicates statistically significant change)

LDH level was lifted more than 4 times of healthy group by the implementation of Carbon Tetrachloride which showed massive contrast with the negative control group. However, treatment groups displayed good results by the huge drop of this LDH level successfully [**Figure 6**]. Same kinds of effects have found in some studies on *Solanum xanthocarpum, Beta vulgaris, Borago officinalis, Aegle marmelos*, and *Mentha piperita* [5], [15], [19–21].

**Figure 6:**
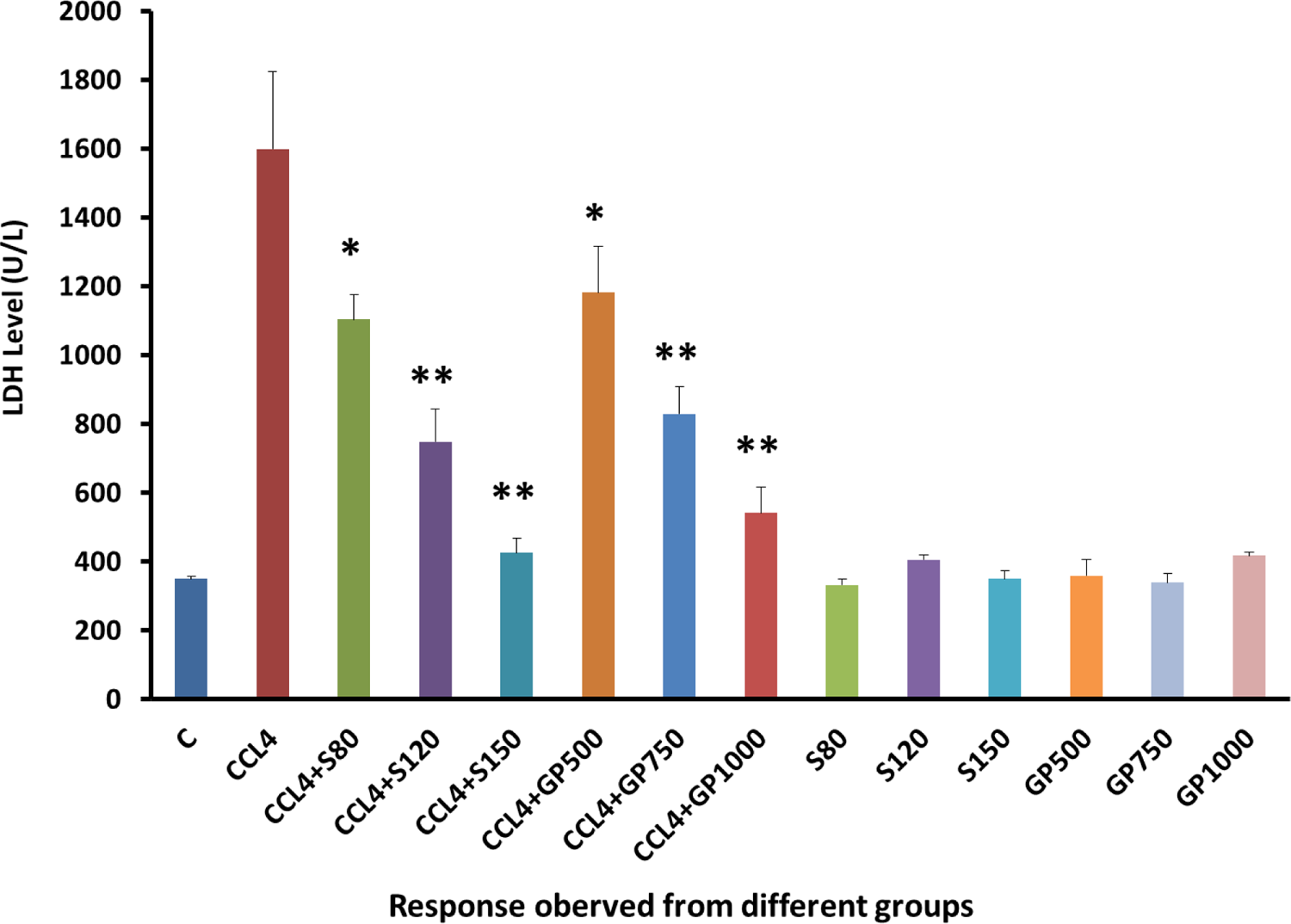
LDH (U/L) Level of rats from 14 groups. The data were expressed as mean ± standard deviation. (* indicates statistically significant change)

In the hepatotoxic specimen, *Gynura procumbens* extract successfully lifted the HDL level in a dose-dependent manner [**Figure 7**]. No observable severe effects were found upon the drug or plant extract administration to the non-CCl_4_ treated groups pointing to the safety of the plant extract. This type of increase in HDL level observed in case of *Nigella sativa, Aegle marmelos,* and *Phytolacca dodecandra* [1], [21], [22]

**Figure 7:**
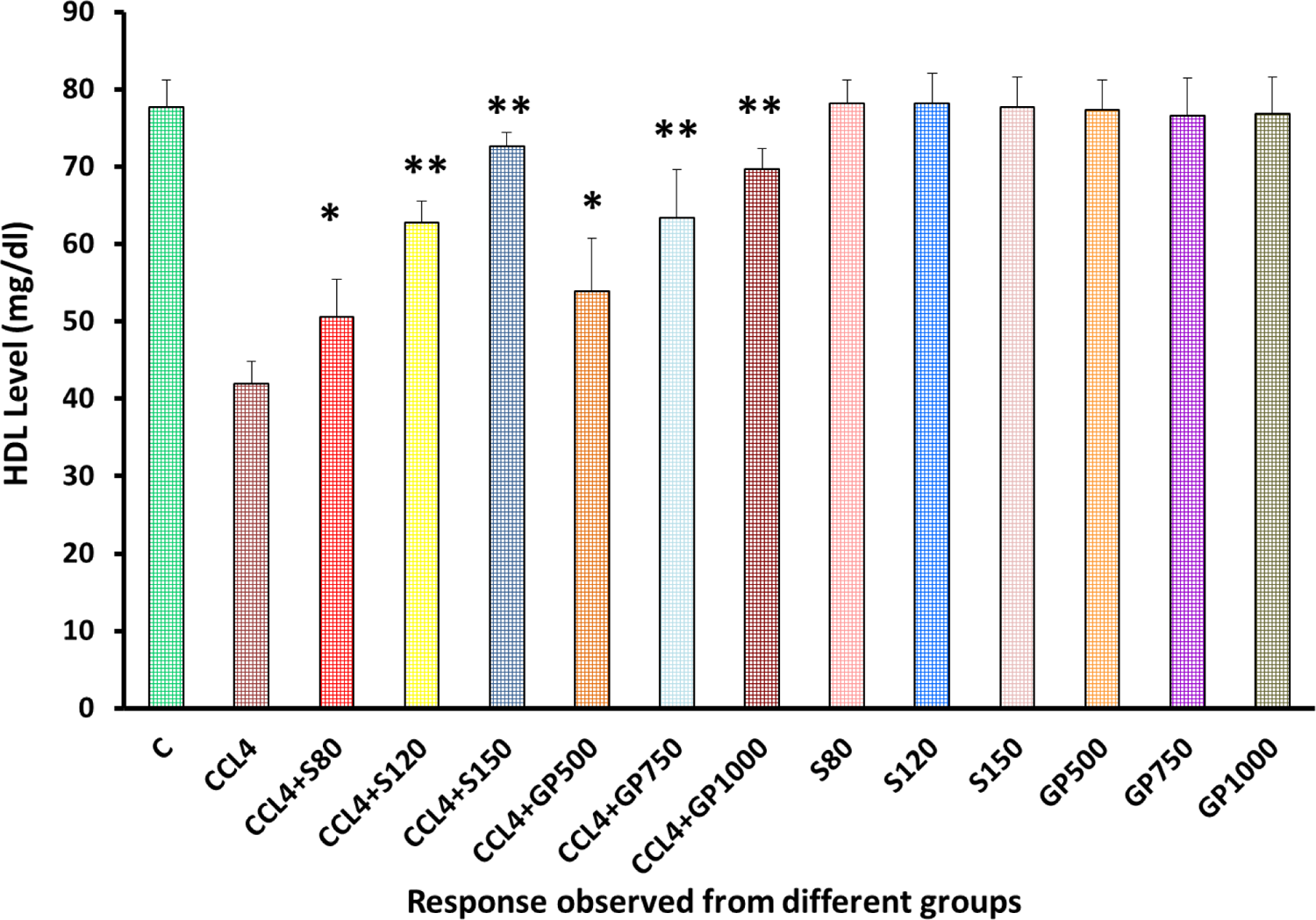
HDL (mg/dl) Level of rats from 14 groups. The data were expressed as mean± standard deviation. (* indicates statistically significant change)

Compare to the negative control group, the disease control group yielded an increasing level of LDL following carbon Tetrachloride administration [**Figure 8**]. Hepatotoxic rats with the treatment of either Silymarin or *Gynura procumbens* extract giving with different doses overturned Carbon Tetrachloride mediated hike in LDL successfully. Lowering of LDL level were also observed in case of *Nigella sativa, Aegle marmelos,* and *Phytolacca dodecandra* [1], [21], [22].

**Figure 8:**
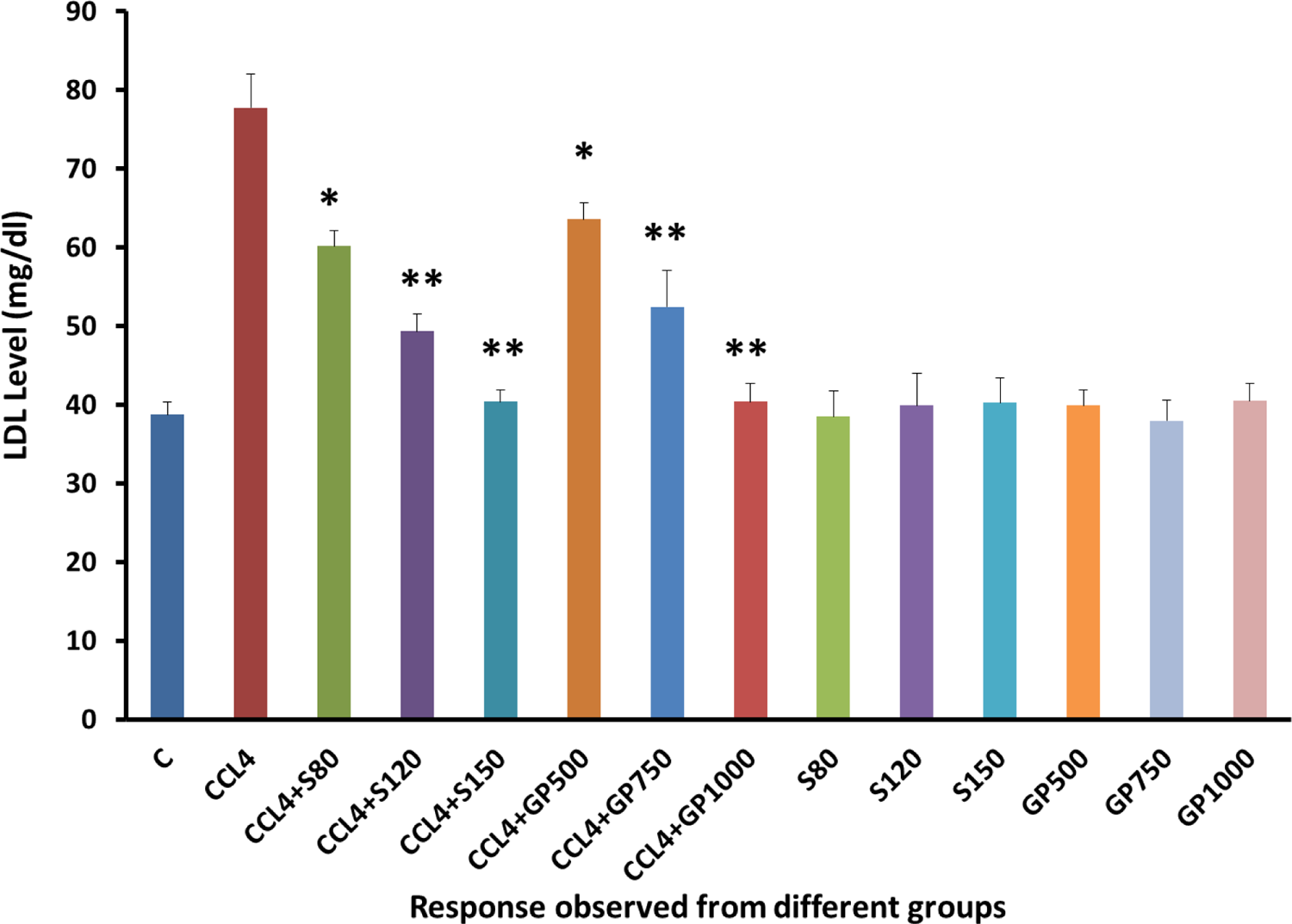
LDL (mg/dl) Level of rats from 14 groups. The data were expressed as mean ± standard deviation. (* indicates statistically significant change)

**Figure 9:**
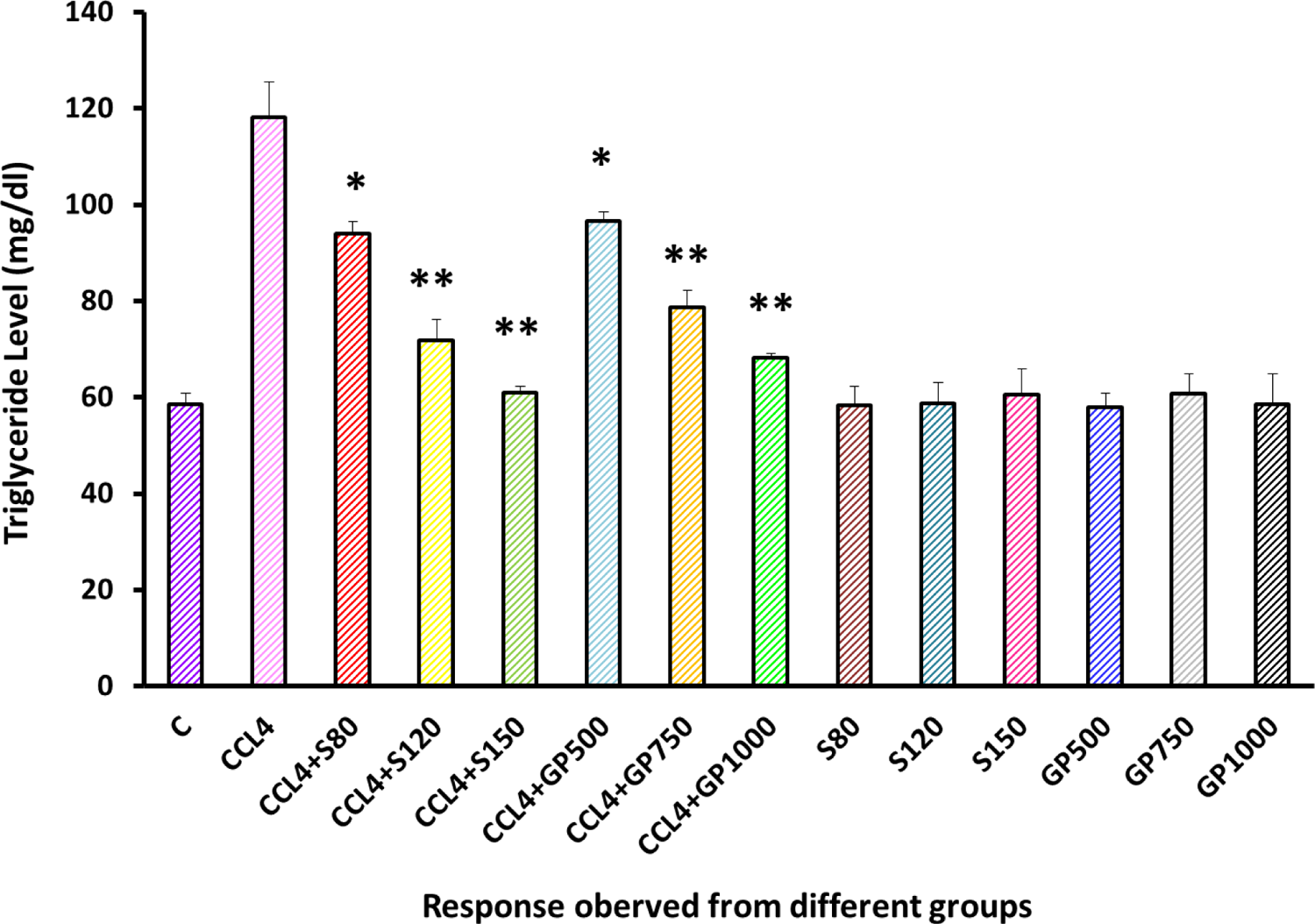
Triglyceride (mg/dl) Level of rats from 14 groups. The data were expressed as mean ± standard deviation. (* indicates statistically significant change)

Triglyceride level was induced almost double due to the application of carbon tetrachloride. However, this level went down to almost same level by the treatment species when highest dose applied. Same kinds of results have obtained for *Clerodendrum inerme,* and *Rhododendron arboreum* plant extracts [4], [14].

Elevated cholesterol level was found in disease control group [**Figure 10**]. Similar pattern was observed both the drugs and extracts where they reduced successfully the cholesterol levels with different doses. Supplementation of highest dose showed the cholesterol level same as the negative control group. There are some studies found to be coincided with our results, using *Beta vulgaris, Clerodendrum inerme, Rhododendron arboreum*, and *Argyreia speciose* plant extracts [2], [4], [5], [14].

**Figure 10:**
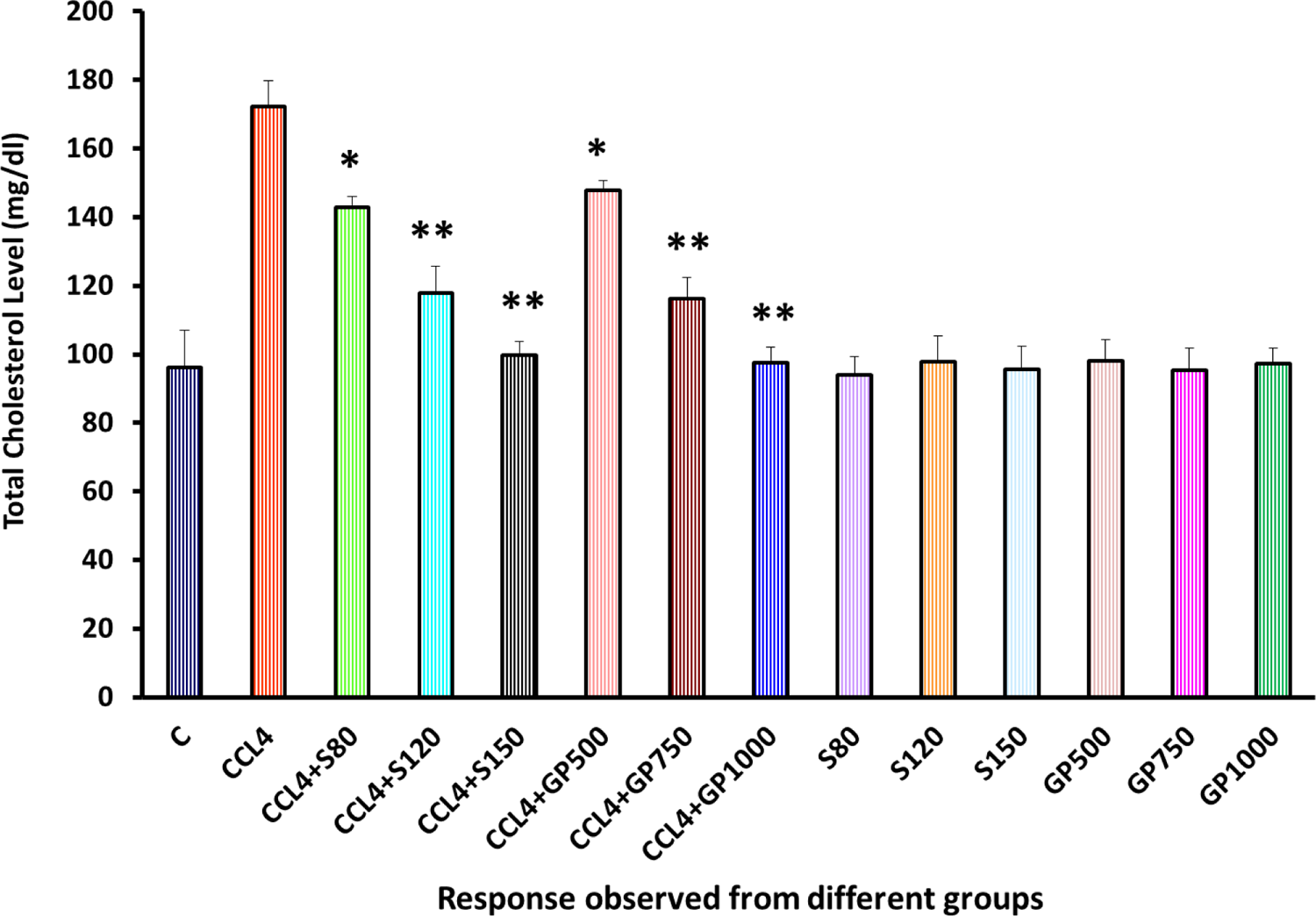
Total Cholesterol (mg/dl) Level of rats from 14 groups. The data were expressed as mean ± standard deviation. (* indicates statistically significant change)

DNA fragmentation ranges were significantly dropped by the silymarin and plant extracts applying to the diseased groups and at the same time there was no significant changes was observed when treatment performed on healthy groups to observe any changes in DNA.

γGT levels were gradually turned to the same level of healthy group with last highest doses both by silymarin and *Gynura procumbens* extracts. The lowering γGT levels were also found in the *Mentha piperita* and *Garcinia kola* plant extracts [19], [23].

A noticeable difference was observed in the CAT level of the positive and disease control groups, as the reduction of the CAT level was prominent after supplying carbon tetrachloride to the rats [**Figure 13**]. On the other hand, contrasting behavior was observed by both the treatment groups with silymarin or the plant extracts where they significantly reversed the CAT level upon carbon tetrachloride towards normal level. Same type of results has found with following plants-*Macrocybe gigantean, Polygonum cuspidatum, Marrubium vulgare*, and *Terminalia arjuna* [3], [16], [17], [24].

In the SOD (Superoxide dismutase) test, disease control group demonstrated a remarkable turn down of this level which indicates the decrease antioxidant defense by the rodents. However, statistically significant improvement was observed by the treatment groups of Silymarin and *Gynura procumbens*. Some of the previous studies complied with the result in case of *Pisonia aculeate, Zanthoxylum armatum, Carya illinoinensis, Solanum xanthocarpum, Saururus chinensis, Macrocybe gigantean, Argyreia speciose, Marrubium vulgare*, and *Terminalia arjuna* [2], [3], [6], [7], [12], [15–17], [25].

A significant overproduction of MDA (Malondialdehyde) was noticed in the diseased group. After administering test extract and the standard drug, a significant statistical difference was detected in the treatment groups as the rate of MDA level was found to be decreased [**Figure 15**]. Studies on *Zanthoxylum armatum, Saururus chinensis, Carya illinoinensis, Solanum xanthocarpum, Aspalathus linearis, Borago officinalis, Macrocybe gigantean, Terminalia arjuna,* and *Phytolacca dodecandra* have showed same type of effects [1], [3], [6], [7], [15], [16], [20], [25], [26].

Of the 76 phytoconstituents of *G. procumbens*, 11 were alkaloids, 32 phenolic compounds, 13 flavonoids, 15 steroidal compounds, and 5 terpenes [51]. The compounds with their respective compound IDs (to be used henceforth on this manuscript) are presented in **Table 2**.

**Table 2:**
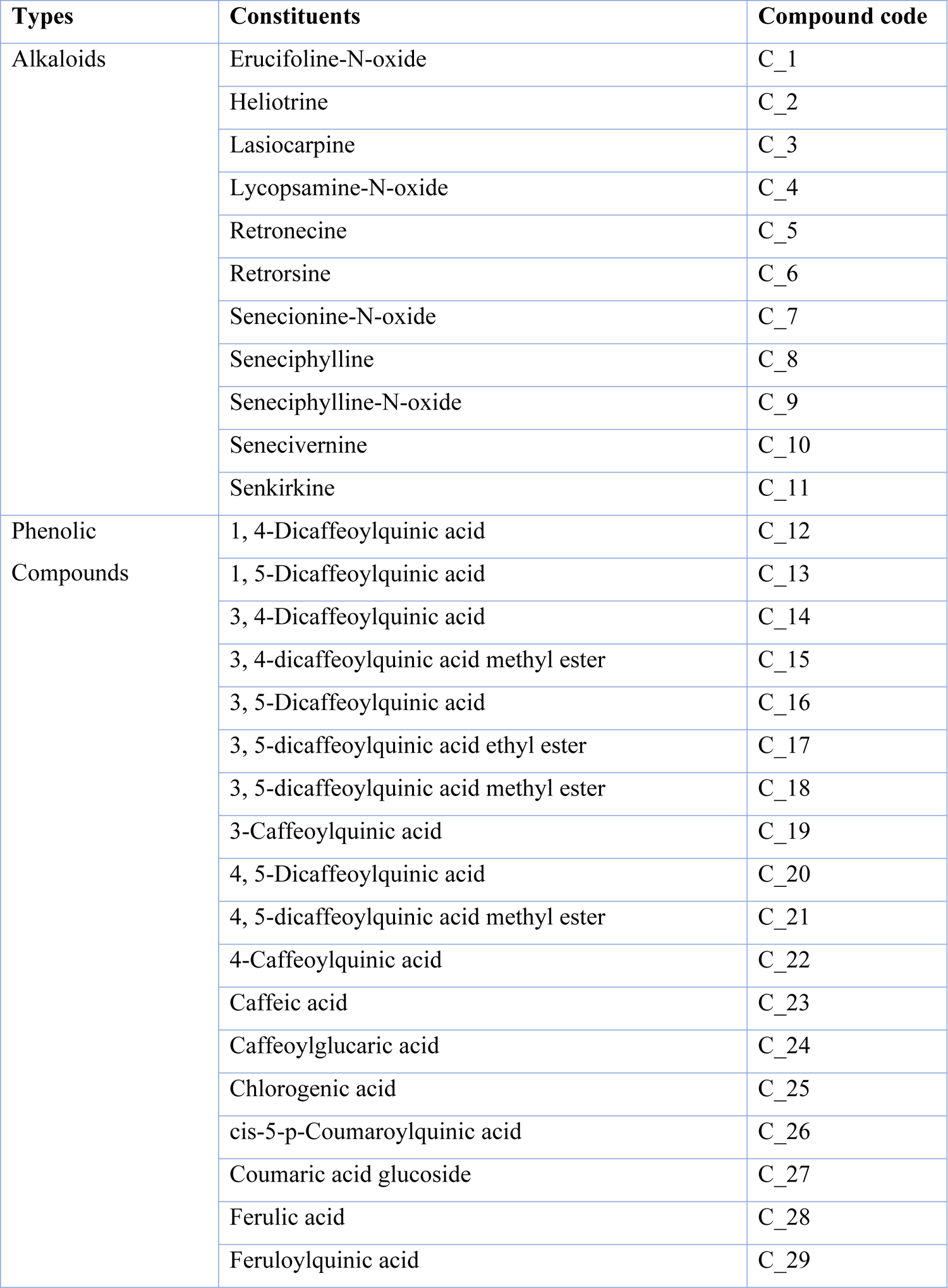
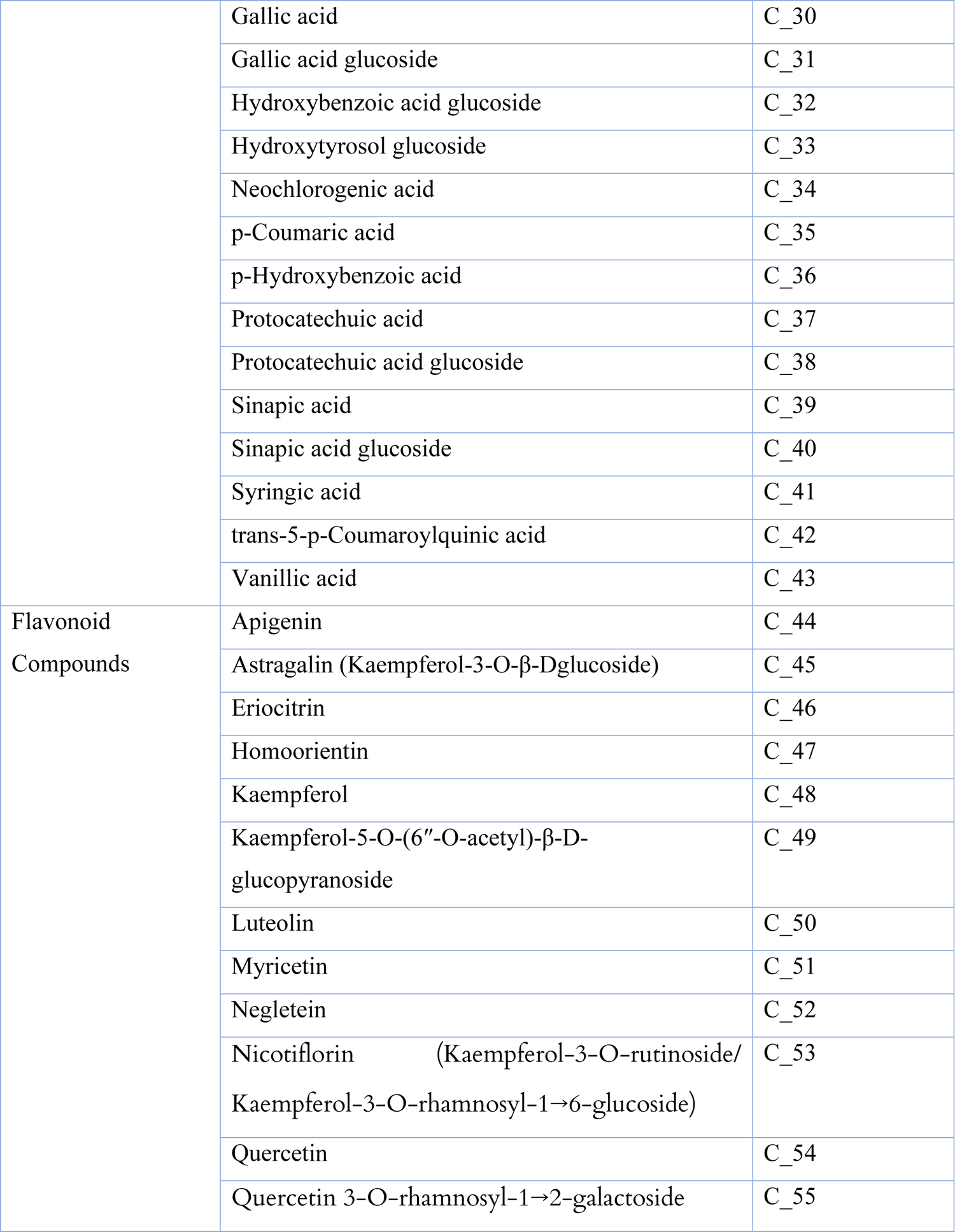
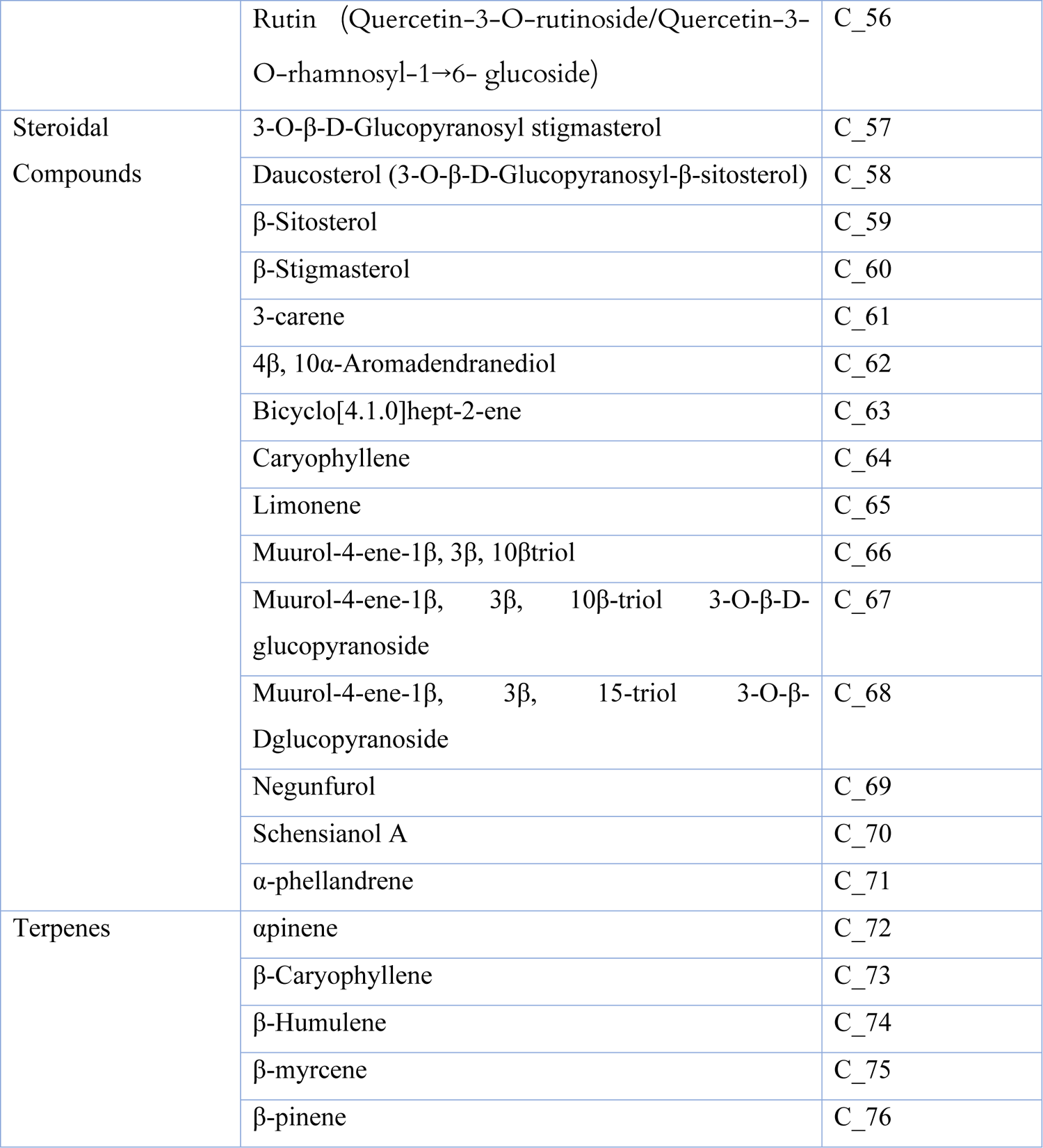
Phytoconstituents of G. procumbens fruit

The macromolecule targets along with their binding sites and respective controls are presented in **Table 3**.

**Table 3:** Macromolecular targets for hepatoprotectivity

| Macromolecule Name | PDB ID | Control drug | Vina Search Space |  | Reference |
| --- | --- | --- | --- | --- | --- |
|  |  |  | Center | Dimensions (Angstrom) |  |
| TGF-β Type I Receptor | 1VJY | Galunisertib | X: 11.8856<br>Y: 66.3307<br>Z: 4.7346 | X: 20.4315<br>Y: 20.1017<br>Z: 18.2275 | 7,8 |
| PPAR-α | 5HYK | Bezafibrate | X: 12.1827<br>Y: 23.8565<br>Z: 25.1561 | X: 17.2809<br>Y: 12.1112<br>Z: 15.6780 | 9,10 |

All 76 phytoconstituents assayed in silico displayed lower binding affinity values than the control Galunisertib (B.A.: −11.5 Kcal/mole) against the TGF-β type I receptor. Compound 50, or luteolin displayed the highest binding affinity after the control at −10.3 Kcal/mole, followed by compound 51 (myricetin) at −10.2 Kcal/mole. Of the top 10 highest binding affinity phytoconstituents, 7 were flavonoids, while the rest were phenolic compounds. All phytoconstituents shared interaction residues with the control. These results are demonstrated in **Table 4**.

**Table 4:** Interactions of phytoconstituents with TGF-β type I receptor

| Ligand | Binding Affinity (Kcal/mole) | Ligand-Macromolecule Interactions |
| --- | --- | --- |
| Galunisertib (control) | -11.5 | ILE A:211*, VAL A:219*, ALA A:230*, LYS A:232*, TYR A:249*, LEU A:260*, LEU A:278*, ASP A:281*, TYR A:282*, HIS A:283*, LEU A:340*, ASP A:351* |
| C_50 | -10.3 | VAL A:219*, ALA A:230*, LYS A:232*, LEU A:260*, LEU A:340*, ALA A:350, ASP A:351* |
| C_51 | -10.2 | VAL A:219*, ALA A:230*, LYS A:232*, GLU A:245, LEU A:260*, LEU A:278*, SER A:280, SER A:287, LEU A:340*, ALA A:350, ASP A:351* |
| C_20 | -10.1 | LYS A:213, ALA A:230*, LYS A:232*, LEU A:260*, HIS A:283*, LEU A:340* |
| C_44 | -10.1 | VAL A:219*, ALA A:230*, LYS A:232*, LEU A:260*, LEU A:278*, SER A:280, SER A:287, LEU A:340*, ALA A:350, ASP A:351* |
| C_48 | -10.1 | VAL A:219*, ALA A:230*, LYS A:232*, LEU A:260*, GLY A:286, SER A:287, LEU A:340*, ALA A:350, ASP A:351* |
| C_13 | -10 | ILE A:211*, GLY A:214, ALA A:230*, LYS A:232*, LEU A:260*, HIS A:283*, SER A:287, LYS A:337, ASP A:351* |
| C_54 | -10 | VAL A:219*, ALA A:230*, LYS A:232*, GLU A:245, LEU A:260*, LEU A:278*, SER A:280, GLY A:286, LEU A:340*, ALA A:350, ASP A:351* |
| C_45 | -9.7 | ILE A:211*, VAL A:219*, ALA A:230*, LYS A:232*, LEU A:260*, TYR A:282*, SER A:287, ASP A:290, LYS A:337, LEU A:340*, ALA A:350, ASP A:351* |
| C_52 | -9.7 | ILE A:211*, VAL A:219*, ALA A:230*, LYS A:232*, LEU A:260*, LEU A:278*, LEU A:340*, ALA A:350, ASP A:351* |
| C_16 | -9.6 | ILE A:211*, VAL A:219*, LYS A:232*, LEU A:260*, HIS A:283*, SER A:287, ASN A:338, LEU A:340* |
| C_14 | -9.5 | ILE A:211*, GLY A:214, VAL A:219*, ALA A:230*, LYS A:232*, LEU A:260*, ASP A:281*, HIS A:283*, ASP A:290, LYS A:335, LEU A:340*, ASN A:338, ALA A:350 |
| C_12 | -9.4 | ILE A:211*, VAL A:219*, ALA A:230*, LYS A:232*, LEU A:260*, ASP A:281*, HIS A:283*, LEU A:340* |
| C_46 | -9.2 | ILE A:211*, GLY A:212, LYS A:213, VAL A:219*, ALA A:230*, LYS A:232*, GLU A:245, LEU A:260*, ASP A:281*, HIS A:283*, SER A:287, LYS A:335, LYS A:337, ASN A:338, LEU A:340*, ALA A:350, ASP A:351* |
| C_17 | -9.1 | LYS A:213, VAL A:219*, ALA A:230*, LYS A:232*, LEU A:260*, LYS A:337, LEU A:340* |
| C_56 | -9.1 | ILE A:211*, LYS A:213, VAL A:219*, ALA A:230*, LYS A:232*, GLU A:245, LEU A:260*, SER A:287, ASP A:290, LYS A:337, LEU A:340*, ALA A:350 |
| C_19 | -8.8 | ILE A:211*, VAL A:219*, LYS A:232*, LEU A:260*, LEU A:278*, SER A:280, SER A:287, LYS A:337 |
| C_34 | -8.8 | VAL A:219*, LYS A:232*, TYR A:249*, LEU A:260*, ALA A:350, ASP A:351* |
| C_15 | -8.6 | ILE A:211*, LYS A:232*, LEU A:260*, SER A:287, LYS A:337 |
| C_33 | -8.6 | LYS A:232*, LEU A:260*, LEU A:278*, HIS A:283* |
| C_22 | -8.5 | LYS A:213, VAL A:219*, ALA A:230*, LYS A:332, LEU A:260*, LYS A:337, ASP A:351* |
| C_25 | -8.5 | VAL A:219*, LYS A:232*, GLU A:245, LEU A:260*, ALA A:350 |
| C_27 | -8.4 | VAL A:219*, ALA A:230*, LEU A:278*, SER A:287, ASN A:338, LEU A:340*, ALA A:350 |
| C_29 | -8.4 | LYS A:213, VAL A:219*, ALA A:230*, LYS A:232*, TYR A:249*, LEU A:260*, PHE A:262, LEU A:278*, SER A:280 |
| C_42 | -8.3 | LYS A:232*, TYR A:249*, LEU A:260*, ASN A:338, ASP A:351* |
| C_49 | -8.3 | LYS A:213, VAL A:219*, ALA A:230*, LYS A:232*, LEU A:260*, SER A:287, LYS A:335, LEU A:340*, ALA A:350, ASP A:351* |
| C_67 | -8.2 | ILE A:211*, GLY A:212, GLY A:214, VAL A:219*, ALA A:230*, LYS A:337, ASN A:338, LEU A:340*, ALA A:350 |
| C_73 | -8.1 | ILE A:211*, VAL A:219*, ALA A:230*, LEU A:260*, TYR A:282*, HIS A:283*, LEU A:340* |
| C_24 | -8 | LYS A:213, VAL A:219*, ALA A:230*, LYS A:232*, LEU A:260*, SER A:280, LYS A:337, ASN A:338, ASP A:351* |
| C_26 | -7.9 | ILE A:211*, VAL A:219*, ALA A:230*, ASP A:281*, HIS A:283*, LEU A:340* |
| C_3 | -7.9 | ILE A:211*, VAL A:219*, ALA A:230*, LYS A:232*, SER A:280, TYR A:282*, LYS A:337, LEU A:340*, ASP A:351* |
| C_1 | -7.8 | GLU A:245, LEU A:340*, ASP A:351* |
| C_55 | -7.8 | ILE A:211*, VAL A:219*, LYS A:232*, GLU A:245, LEU A:278*, SER A:280, TYR A:282*, HIS A:283*, HIS A:285, GLY A:286, LYS A:337, LEU A:340*, ASP A:351* |
| C_6 | -7.8 | ILE A:211*, VAL A:219*, ASP A:281*, SER A:287, LEU A:340* |
| C_60 | -7.8 | ILE A:211*, VAL A:219*, ALA A:230*, LYS A:232*, LEU A:260*, LYS A:337, LEU A:340*, ALA A:350 |
| C_32 | -7.7 | LYS A:213, VAL A:219*, ALA A:230*, LYS A:232*, LEU A:260*, LEU A:278*, ALA A:350, ASP A:351* |
| C_40 | -7.7 | VAL A:219*, ALA A:230*, LEU A:278*, SER A:287, ASP A:290, LEU A:340*, ALA A:350, ASP A:351* |
| C_64 | -7.7 | ILE A:211*, VAL A:219*, ALA A:230*, LEU A:260*, LEU A:340*, ALA A:350 |
| C_68 | -7.7 | ILE A:211*, LYS A:213, GLY A:214, VAL A:219*, ALA A:230*, LYS A:337, ASN A:338, LEU A:340*, ASP A:351* |
| C_7 | -7.7 | ILE A:211*, SER A:287, LEU A:340*, ASP A:351* |
| C_11 | -7.6 | ILE A:211*, VAL A:219*, ALA A:230*, TYR A:282*, SER A:287, ASN A:338, LEU A:340* |
| C_31 | -7.6 | VAL A:219*, ALA A:230*, LYS A:232*, LEU A:260*, LEU A:278*, ALA A:350, ASP A:351* |
| C_38 | -7.6 | ILE A:211*, GLY A:214, VAL A:219*, ALA A:230*, HIS A:283*, SER A:287, ASN A:338, LEU A:340*, ASP A:351* |
| C_18 | -7.5 | LYS A:213, GLY A:214, ALA A:230*, LYS A:232*, LEU A:260*, LEU A:278*, SER A:280, ASP A:281*, HIS A:283*, LYS A:337, ASN A:338, LEU A:340* |
| C_9 | -7.4 | ILE A:211*, VAL A:219*, ALA A:230*, LEU A:260*, SER A:287, LYS A:337, LEU A:340*, ASP A:351* |
| C_62 | -7.3 | ILE A:211*, VAL A:219*, ALA A:230*, HIS A:283*, LEU A:340*, ALA A:350 |
| C_66 | -7.3 | ILE A:211*, VAL A:219*, LEU A:260*, LEU A:340*, ALA A:350, ASP A:351* |
| C_69 | -7.2 | VAL A:219*, ALA A:230*, LYS A:232*, TYR A:249*, LEU A:260*, PHE A:262, LEU A:278*, HIS A:283*, LEU A:340*, ALA A:350 |
| C_70 | -7.2 | VAL A:219*, ALA A:230*, LYS A:232*, TYR A:249*, LEU A:260*, PHE A:262, LEU A:278*, LEU A:340*, ALA A:350 |
| C_74 | -7.2 | ILE A:211*, VAL A:219*, ALA A:230*, LEU A:260*, TYR A:282*, HIS A:283*, LEU A:340* |
| C_2 | -7.1 | ILE A:211*, VAL A:219*, ALA A:230*, LYS A:232*, TYR A:249*, LEU A:260*, LEU A:278*, ASP A:351* |
| C_59 | -7.1 | ILE A:211*, LYS A:213, VAL A:219*, ALA A:230*, LYS A:232*, LEU A:260*, TYR A:282*, LYS A:337, LEU A:340*, ALA A:350 |
| C_21 | -7 | VAL A:219*, ALA A:230*, LYS A:232*, GLU A:245, LEU A:260*, HIS A:283*, GLY A:286, SER A:287, ASP A:290, LYS A:337, ASN A:338, ALA A:350, ASP A:351* |
| C_23 | -6.9 | VAL A:219*, ALA A:230*, LYS A:232*, ASP A:281*, HIS A:283*, LEU A:340*, ASP A:351* |
| C_37 | -6.9 | VAL A:219*, ALA A:230*, LYS A:232*, GLU A:245, TYR A:249*, LEU A:260* |
| C_28 | -6.8 | VAL A:219*, ALA A:230*, LYS A:232*, LEU A:340* |
| C_36 | -6.8 | VAL A:219*, ALA A:230*, LYS A:232*, LEU A:260* |
| C_39 | -6.8 | VAL A:219*, ALA A:230*, LYS A:232*, LEU A:260*, LYS A:337, LEU A:340*, ALA A:350, ASP A:351* |
| C_30 | -6.7 | ALA A:230*, LYS A:232*, LEU A:260*, SER A:280, ASP A:351* |
| C_43 | -6.7 | VAL A:219*, ALA A:230*, LYS A:232*, TYR A:249*, LEU A:260*, PHE A:262, LEU A:278*, SER A:280, ASP A:351* |
| C_35 | -6.6 | VAL A:219*, ALA A:230*, LYS A:232*, HIS A:283*, LEU A:340*, ASP A:351* |
| C_4 | -6.6 | VAL A:219*, LYS A:232*, SER A:280, LYS A:337, ASN A:338, ASP A:351* |
| C_57 | -6.5 | ILE A:211*, VAL A:219*, ALA A:230*, LYS A:232*, LEU A:260*, HIS A:283*, LYS A:337, LEU A:340*, ALA A:350 |
| C_71 | -6.3 | VAL A:219*, LYS A:232*, TYR A:249*, LEU A:260*, LEU A:278*, LEU A:340*, ALA A:350 |
| C_10 | -6.2 | VAL A:219*, ALA A:230*, LEU A:260*, LYS A:337, LEU A:340*, ALA A:350 |
| C_47 | -6.2 | VAL A:219*, ALA A:230*, LYS A:232*, SER A:287, ASP A:290, LYS A:337, LEU A:340*, ALA A:350 |
| C_65 | -6.1 | ILE A:211*, VAL A:219*, ALA A:230*, LEU A:260*, TYR A:282*, LEU A:340* |
| C_41 | -6 | ILE A:211*, VAL A:219*, ALA A:230*, LYS A:232*, LEU A:260*, HIS A:283*, LEU A:340* |
| C_8 | -5.9 | ILE A:211*, VAL A:219*, ALA A:230*, LYS A:232*, LEU A:340*, ALA A:350 |
| C_5 | -5.7 | VAL A:219*, ALA A:230*, LYS A:232*, GLU A:245, TYR A:249*, LEU A:260*, LEU A:278*, ASP A:351* |
| C_75 | -5.7 | VAL A:219*, ALA A:230*, LYS A:232*, TYR A:249*, LEU A:260*, PHE A:262, LEU A:278*, LEU A:340*, ALA A:350 |
| C_61 | -5.6 | ILE A:211*, VAL A:219*, LEU A:340*, ALA A:350 |
| C_53 | -5.2 | ILE A:211*, VAL A:219*, ALA A:230*, LYS A:232*, LEU A:260*, SER A:287, LYS A:337, ASN A:338, LEU A:340*, ALA A:350, ASP A:351* |
| C_63 | -5.1 | LYS A:232*, TYR A:249*, LEU A:260*, PHE A:262, LEU A:278* |
| C_76 | -5 | VAL A:219*, ALA A:230*, LYS A:232*, LEU A:260*, LEU A:340*, ALA A:350 |
| C_72 | -4.8 | VAL A:219*, ALA A:230*, LYS A:232*, LEU A:260*, LEU A:340*, ALA A:350 |
| C_58 | 1.1 | VAL A:219*, ALA A:230*, LYS A:232*, LEU A:260*, LEU A:278*, SER A:280, LYS A:337, LEU A:340*, ALA A:350 |

Interactions common with the control\*

26 phytoconstituents out of 76 displayed higher binding affinity values than the control bezafibrate (B.A.: −7.6 Kcal/mole) against PPAR-α. Compound 8, or seneciphylline displayed the highest binding affinity after the control at −8.7 Kcal/mole, followed by compound 26 (cis-5-p-coumaroylquinic acid) at −8.6 Kcal/mole and compound 27 (coumaric acid glucoside) at −8.5 Kcal/mole. Of these 26, 15 were phenolic compounds, 6 were flavonoids, 2 were steroidal compounds, 2 were terpenes, and 1 was an alkaloid. Of the top 10 highest binding affinity phytoconstituents, 1 was an alkaloid, 6 were flavonoids, and 3 were phenolic compounds. All phytoconstituents except compounds 25, 26, 34, and 38 shared interaction residues with the control. These results are demonstrated in **Table 5**.

**Table 5:**
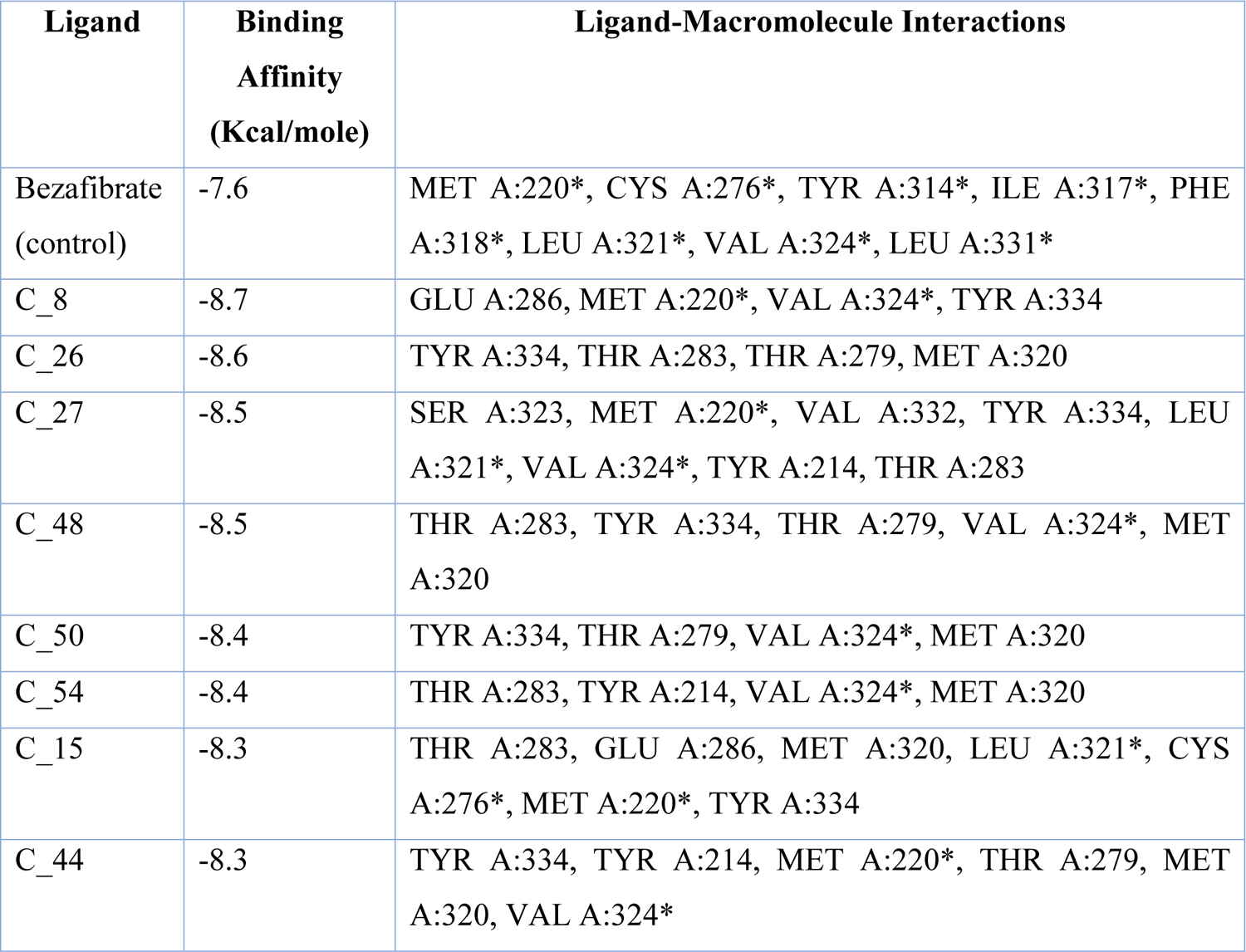
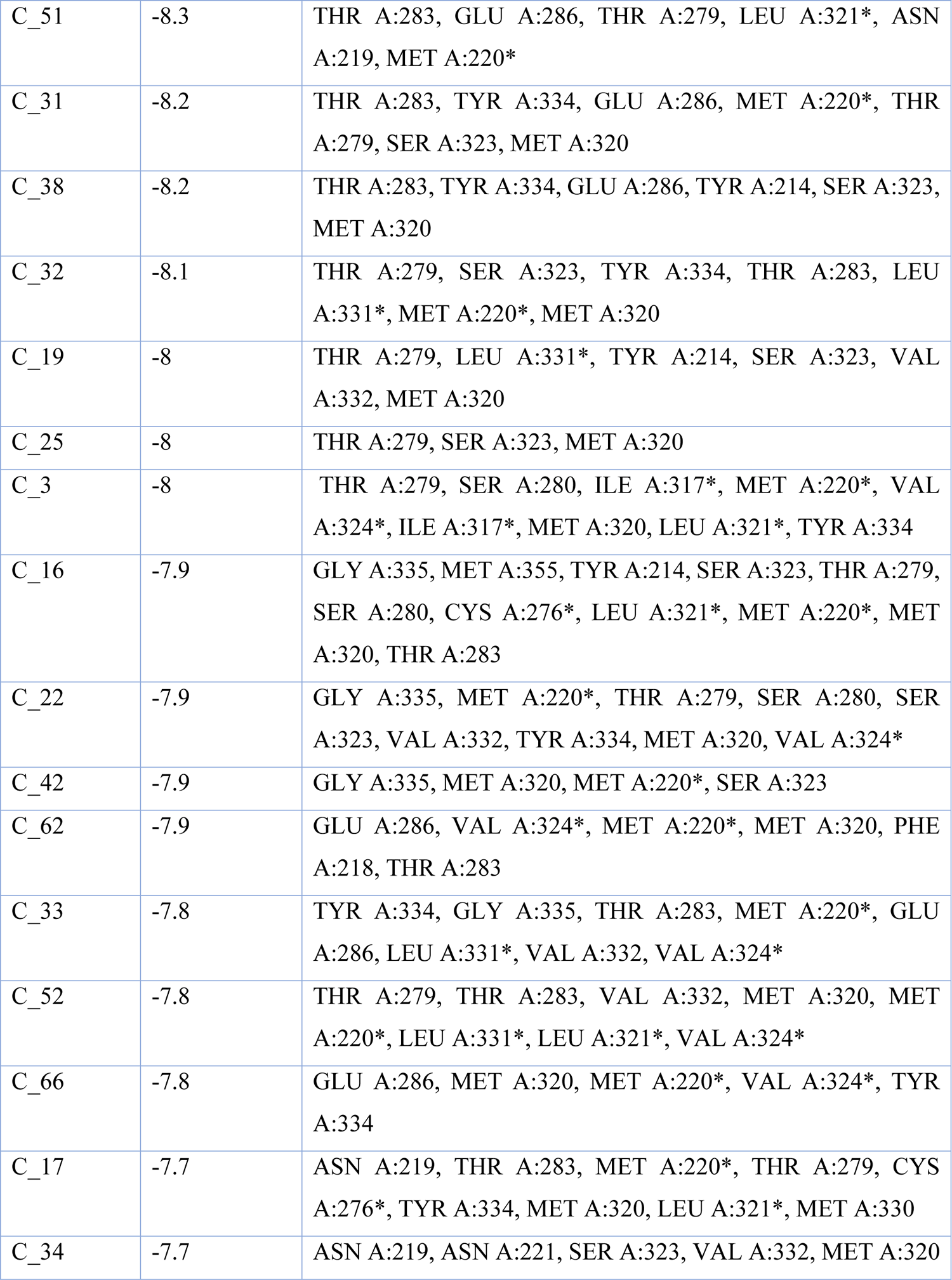
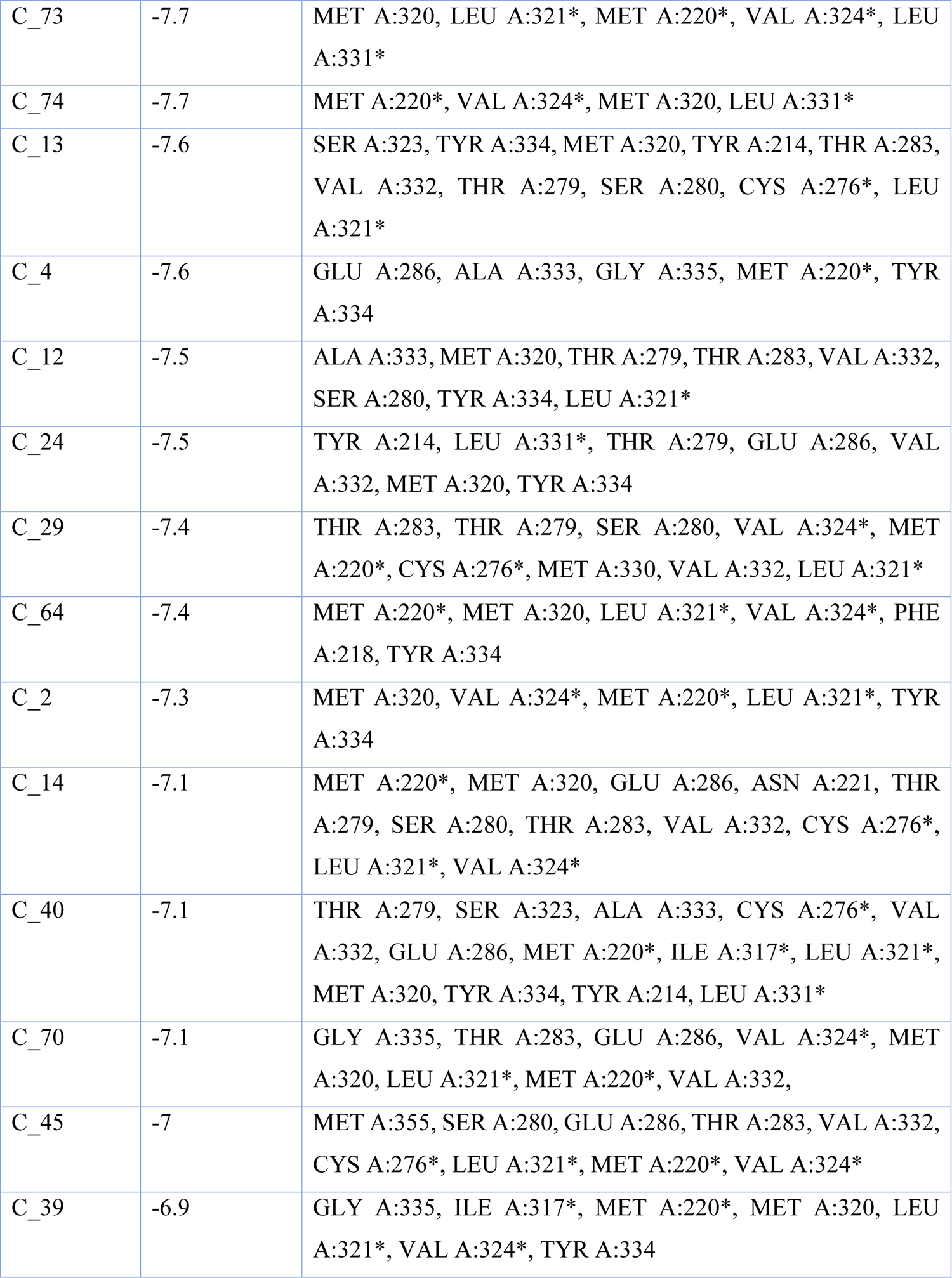
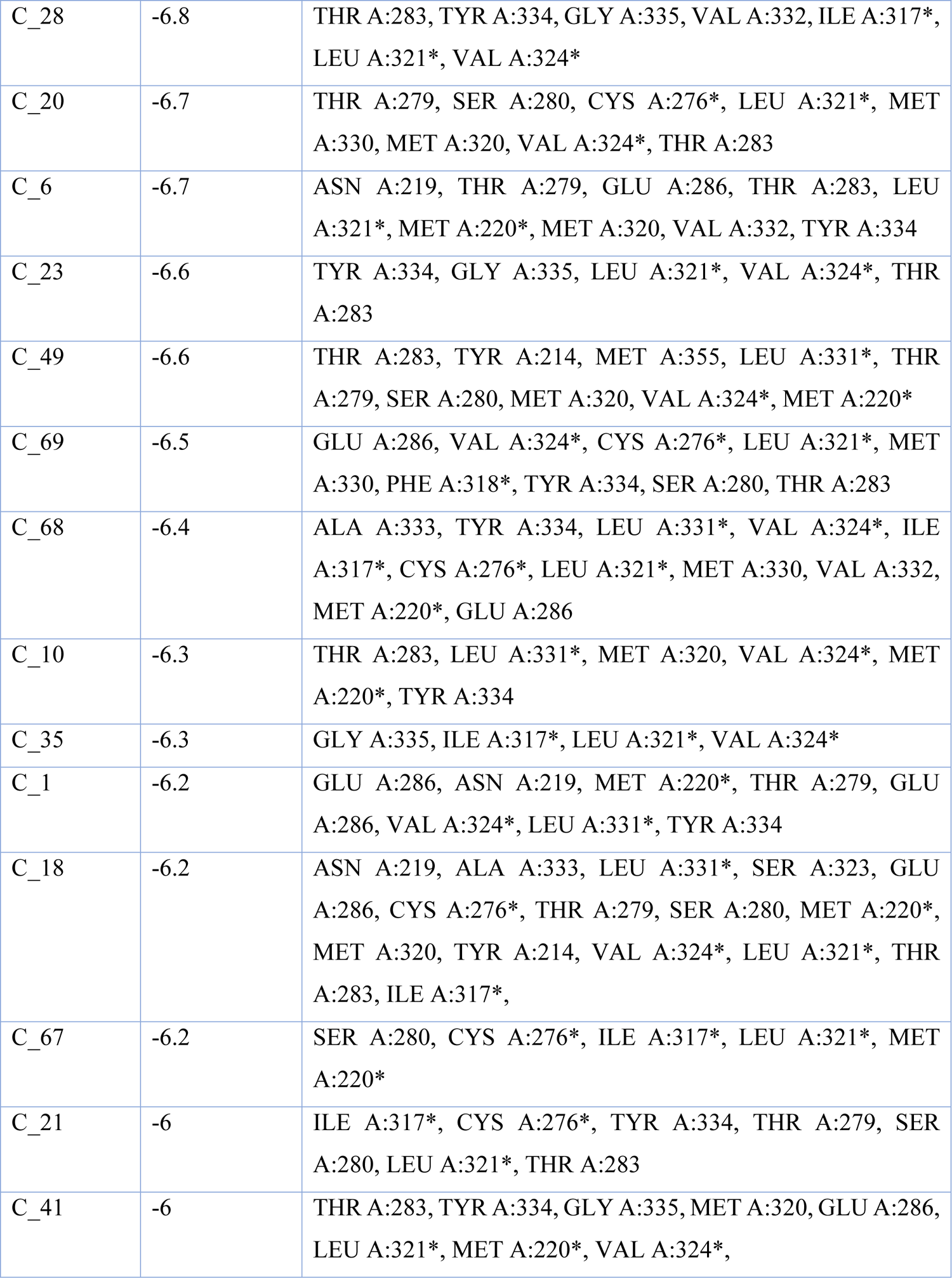
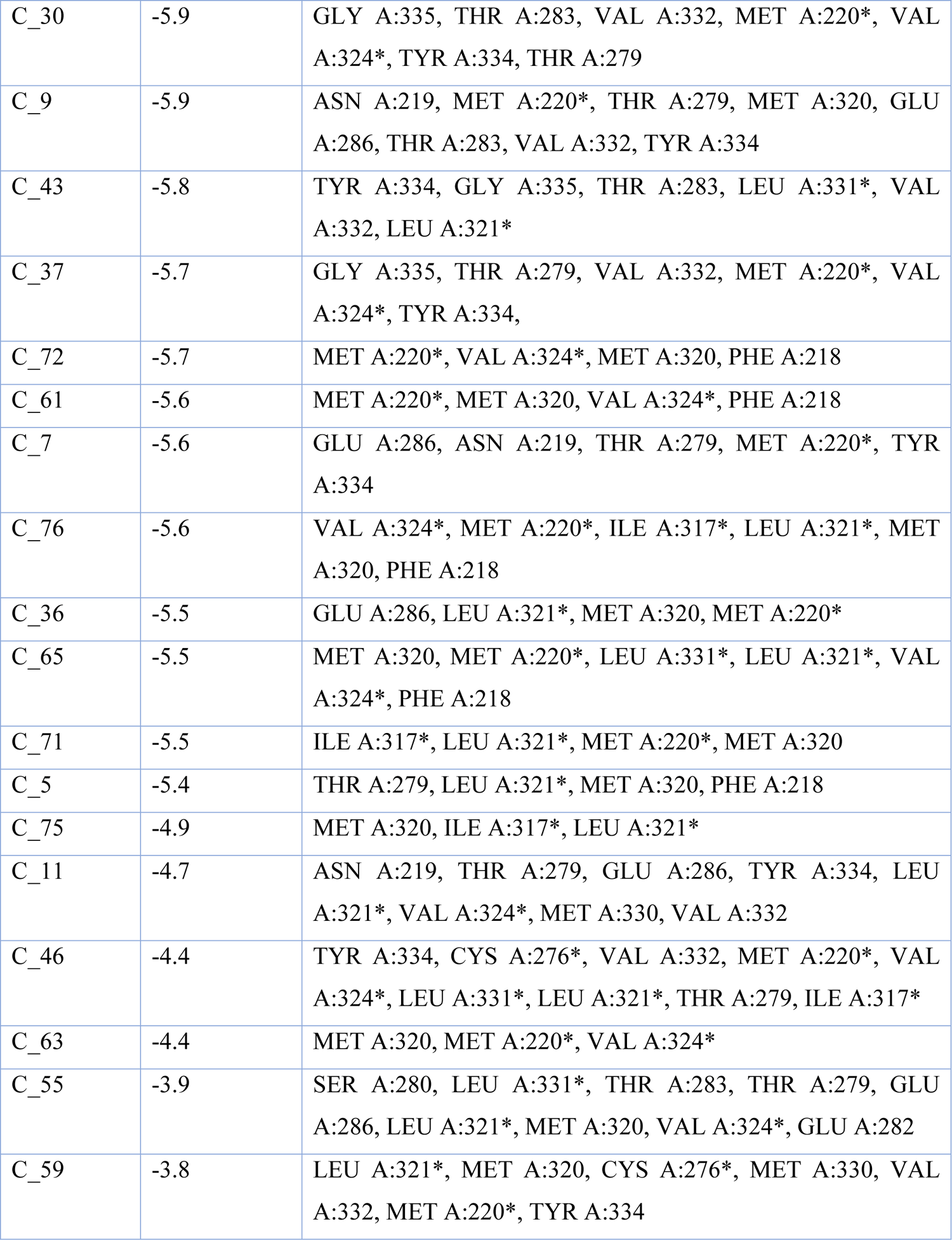
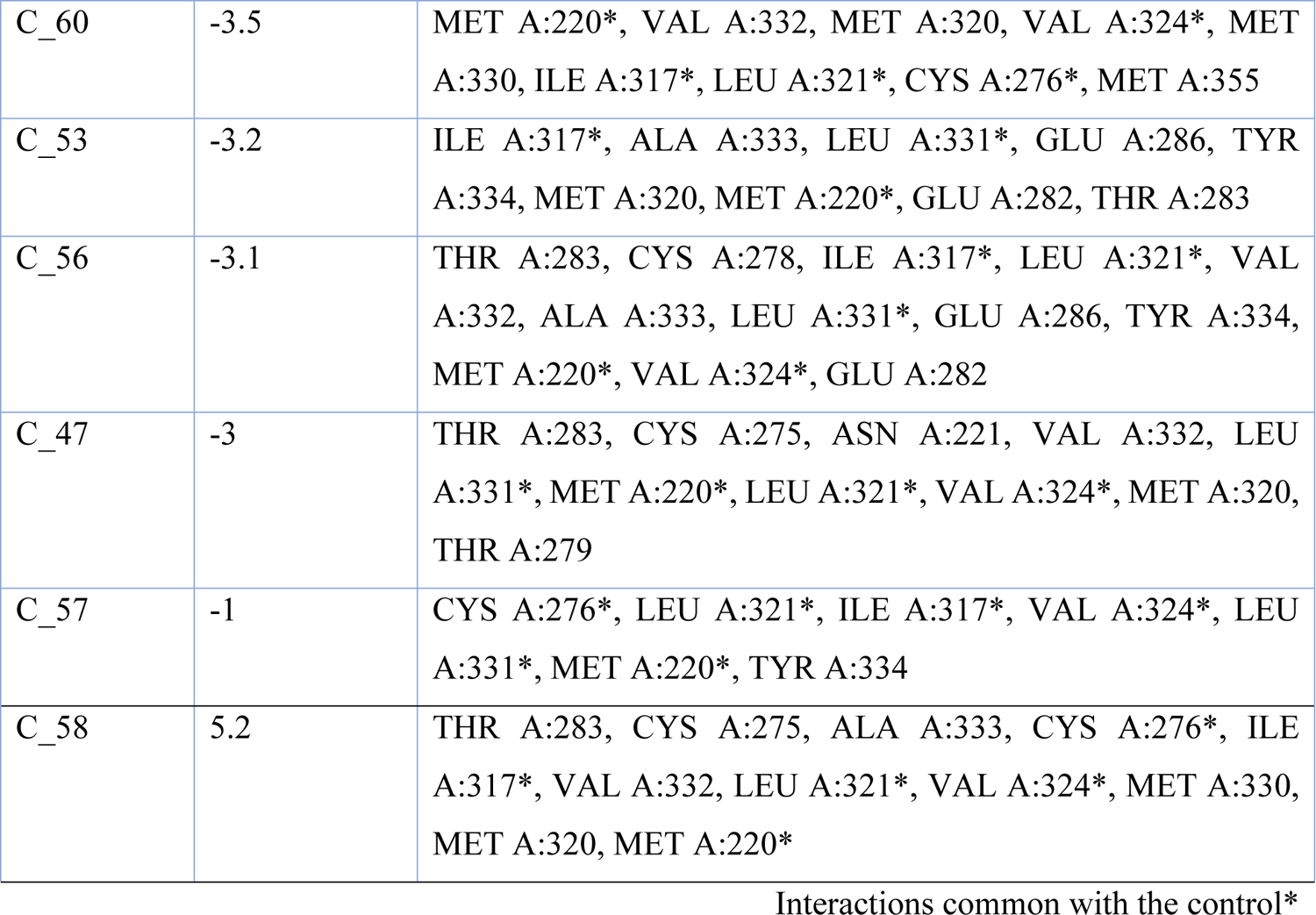
Interactions of phytoconstituents with PPAR-α

| Ligand | Binding Affinity (Kcal/mole) | Ligand-Macromolecule Interactions |
| --- | --- | --- |
| Bezafibrate (control) | -7.6 | MET A:220*, CYS A:276*, TYR A:314*, ILE A:317*, PHE A:318*, LEU A:321*, VAL A:324*, LEU A:331* |
| C_8 | -8.7 | GLU A:286, MET A:220*, VAL A:324*, TYR A:334 |
| C_26 | -8.6 | TYR A:334, THR A:283, THR A:279, MET A:320 |
| C_27 | -8.5 | SER A:323, MET A:220*, VAL A:332, TYR A:334, LEU A:321*, VAL A:324*, TYR A:214, THR A:283 |
| C_48 | -8.5 | THR A:283, TYR A:334, THR A:279, VAL A:324*, MET A:320 |
| C_50 | -8.4 | TYR A:334, THR A:279, VAL A:324*, MET A:320 |
| C_54 | -8.4 | THR A:283, TYR A:214, VAL A:324*, MET A:320 |
| C_15 | -8.3 | THR A:283, GLU A:286, MET A:320, LEU A:321*, CYS A:276*, MET A:220*, TYR A:334 |
| C_44 | -8.3 | TYR A:334, TYR A:214, MET A:220*, THR A:279, MET A:320, VAL A:324* |
| C_51 | -8.3 | THR A:283, GLU A:286, THR A:279, LEU A:321*, ASN A:219, MET A:220* |
| C_31 | -8.2 | THR A:283, TYR A:334, GLU A:286, MET A:220*, THR A:279, SER A:323, MET A:320 |
| C_38 | -8.2 | THR A:283, TYR A:334, GLU A:286, TYR A:214, SER A:323, MET A:320 |
| C_32 | -8.1 | THR A:279, SER A:323, TYR A:334, THR A:283, LEU A:331*, MET A:220*, MET A:320 |
| C_19 | -8 | THR A:279, LEU A:331*, TYR A:214, SER A:323, VAL A:332, MET A:320 |
| C_25 | -8 | THR A:279, SER A:323, MET A:320 |
| C_3 | -8 | THR A:279, SER A:280, ILE A:317*, MET A:220*, VAL A:324*, ILE A:317*, MET A:320, LEU A:321*, TYR A:334 |
| C_16 | -7.9 | GLY A:335, MET A:355, TYR A:214, SER A:323, THR A:279, SER A:280, CYS A:276*, LEU A:321*, MET A:220*, MET A:320, THR A:283 |
| C_22 | -7.9 | GLY A:335, MET A:220*, THR A:279, SER A:280, SER A:323, VAL A:332, TYR A:334, MET A:320, VAL A:324* |
| C_42 | -7.9 | GLY A:335, MET A:320, MET A:220*, SER A:323 |
| C_62 | -7.9 | GLU A:286, VAL A:324*, MET A:220*, MET A:320, PHE A:218, THR A:283 |
| C_33 | -7.8 | TYR A:334, GLY A:335, THR A:283, MET A:220*, GLU A:286, LEU A:331*, VAL A:332, VAL A:324* |
| C_52 | -7.8 | THR A:279, THR A:283, VAL A:332, MET A:320, MET A:220*, LEU A:331*, LEU A:321*, VAL A:324* |
| C_66 | -7.8 | GLU A:286, MET A:320, MET A:220*, VAL A:324*, TYR A:334 |
| C_17 | -7.7 | ASN A:219, THR A:283, MET A:220*, THR A:279, CYS A:276*, TYR A:334, MET A:320, LEU A:321*, MET A:330 |
| C_34 | -7.7 | ASN A:219, ASN A:221, SER A:323, VAL A:332, MET A:320 |
| C_73 | -7.7 | MET A:320, LEU A:321*, MET A:220*, VAL A:324*, LEU A:331* |
| C_74 | -7.7 | MET A:220*, VAL A:324*, MET A:320, LEU A:331* |
| C_13 | -7.6 | SER A:323, TYR A:334, MET A:320, TYR A:214, THR A:283, VAL A:332, THR A:279, SER A:280, CYS A:276*, LEU A:321* |
| C_4 | -7.6 | GLU A:286, ALA A:333, GLY A:335, MET A:220*, TYR A:334 |
| C_12 | -7.5 | ALA A:333, MET A:320, THR A:279, THR A:283, VAL A:332, SER A:280, TYR A:334, LEU A:321* |
| C_24 | -7.5 | TYR A:214, LEU A:331*, THR A:279, GLU A:286, VAL A:332, MET A:320, TYR A:334 |
| C_29 | -7.4 | THR A:283, THR A:279, SER A:280, VAL A:324*, MET A:220*, CYS A:276*, MET A:330, VAL A:332, LEU A:321* |
| C_64 | -7.4 | MET A:220*, MET A:320, LEU A:321*, VAL A:324*, PHE A:218, TYR A:334 |
| C_2 | -7.3 | MET A:320, VAL A:324*, MET A:220*, LEU A:321*, TYR A:334 |
| C_14 | -7.1 | MET A:220*, MET A:320, GLU A:286, ASN A:221, THR A:279, SER A:280, THR A:283, VAL A:332, CYS A:276*, LEU A:321*, VAL A:324* |
| C_40 | -7.1 | THR A:279, SER A:323, ALA A:333, CYS A:276*, VAL A:332, GLU A:286, MET A:220*, ILE A:317*, LEU A:321*, MET A:320, TYR A:334, TYR A:214, LEU A:331* |
| C_70 | -7.1 | GLY A:335, THR A:283, GLU A:286, VAL A:324*, MET A:320, LEU A:321*, MET A:220*, VAL A:332, |
| C_45 | -7 | MET A:355, SER A:280, GLU A:286, THR A:283, VAL A:332, CYS A:276*, LEU A:321*, MET A:220*, VAL A:324* |
| C_39 | -6.9 | GLY A:335, ILE A:317*, MET A:220*, MET A:320, LEU A:321*, VAL A:324*, TYR A:334 |
| C_28 | -6.8 | THR A:283, TYR A:334, GLY A:335, VAL A:332, ILE A:317*, LEU A:321*, VAL A:324* |
| C_20 | -6.7 | THR A:279, SER A:280, CYS A:276*, LEU A:321*, MET A:330, MET A:320, VAL A:324*, THR A:283 |
| C_6 | -6.7 | ASN A:219, THR A:279, GLU A:286, THR A:283, LEU A:321*, MET A:220*, MET A:320, VAL A:332, TYR A:334 |
| C_23 | -6.6 | TYR A:334, GLY A:335, LEU A:321*, VAL A:324*, THR A:283 |
| C_49 | -6.6 | THR A:283, TYR A:214, MET A:355, LEU A:331*, THR A:279, SER A:280, MET A:320, VAL A:324*, MET A:220* |
| C_69 | -6.5 | GLU A:286, VAL A:324*, CYS A:276*, LEU A:321*, MET A:330, PHE A:318*, TYR A:334, SER A:280, THR A:283 |
| C_68 | -6.4 | ALA A:333, TYR A:334, LEU A:331*, VAL A:324*, ILE A:317*, CYS A:276*, LEU A:321*, MET A:330, VAL A:332, MET A:220*, GLU A:286 |
| C_10 | -6.3 | THR A:283, LEU A:331*, MET A:320, VAL A:324*, MET A:220*, TYR A:334 |
| C_35 | -6.3 | GLY A:335, ILE A:317*, LEU A:321*, VAL A:324* |
| C_1 | -6.2 | GLU A:286, ASN A:219, MET A:220*, THR A:279, GLU A:286, VAL A:324*, LEU A:331*, TYR A:334 |
| C_18 | -6.2 | ASN A:219, ALA A:333, LEU A:331*, SER A:323, GLU A:286, CYS A:276*, THR A:279, SER A:280, MET A:220*, MET A:320, TYR A:214, VAL A:324*, LEU A:321*, THR A:283, ILE A:317*, |
| C_67 | -6.2 | SER A:280, CYS A:276*, ILE A:317*, LEU A:321*, MET A:220* |
| C_21 | -6 | ILE A:317*, CYS A:276*, TYR A:334, THR A:279, SER A:280, LEU A:321*, THR A:283 |
| C_41 | -6 | THR A:283, TYR A:334, GLY A:335, MET A:320, GLU A:286, LEU A:321*, MET A:220*, VAL A:324*, |
| C_30 | -5.9 | GLY A:335, THR A:283, VAL A:332, MET A:220*, VAL A:324*, TYR A:334, THR A:279 |
| C_9 | -5.9 | ASN A:219, MET A:220*, THR A:279, MET A:320, GLU A:286, THR A:283, VAL A:332, TYR A:334 |
| C_43 | -5.8 | TYR A:334, GLY A:335, THR A:283, LEU A:331*, VAL A:332, LEU A:321* |
| C_37 | -5.7 | GLY A:335, THR A:279, VAL A:332, MET A:220*, VAL A:324*, TYR A:334, |
| C_72 | -5.7 | MET A:220*, VAL A:324*, MET A:320, PHE A:218 |
| C_61 | -5.6 | MET A:220*, MET A:320, VAL A:324*, PHE A:218 |
| C_7 | -5.6 | GLU A:286, ASN A:219, THR A:279, MET A:220*, TYR A:334 |
| C_76 | -5.6 | VAL A:324*, MET A:220*, ILE A:317*, LEU A:321*, MET A:320, PHE A:218 |
| C_36 | -5.5 | GLU A:286, LEU A:321*, MET A:320, MET A:220* |
| C_65 | -5.5 | MET A:320, MET A:220*, LEU A:331*, LEU A:321*, VAL A:324*, PHE A:218 |
| C_71 | -5.5 | ILE A:317*, LEU A:321*, MET A:220*, MET A:320 |
| C_5 | -5.4 | THR A:279, LEU A:321*, MET A:320, PHE A:218 |
| C_75 | -4.9 | MET A:320, ILE A:317*, LEU A:321* |
| C_11 | -4.7 | ASN A:219, THR A:279, GLU A:286, TYR A:334, LEU A:321*, VAL A:324*, MET A:330, VAL A:332 |
| C_46 | -4.4 | TYR A:334, CYS A:276*, VAL A:332, MET A:220*, VAL A:324*, LEU A:331*, LEU A:321*, THR A:279, ILE A:317* |
| C_63 | -4.4 | MET A:320, MET A:220*, VAL A:324* |
| C_55 | -3.9 | SER A:280, LEU A:331*, THR A:283, THR A:279, GLU A:286, LEU A:321*, MET A:320, VAL A:324*, GLU A:282 |
| C_59 | -3.8 | LEU A:321*, MET A:320, CYS A:276*, MET A:330, VAL A:332, MET A:220*, TYR A:334 |
| C_60 | -3.5 | MET A:220*, VAL A:332, MET A:320, VAL A:324*, MET A:330, ILE A:317*, LEU A:321*, CYS A:276*, MET A:355 |
| C_53 | -3.2 | ILE A:317*, ALA A:333, LEU A:331*, GLU A:286, TYR A:334, MET A:320, MET A:220*, GLU A:282, THR A:283 |
| C_56 | -3.1 | THR A:283, CYS A:278, ILE A:317*, LEU A:321*, VAL A:332, ALA A:333, LEU A:331*, GLU A:286, TYR A:334, MET A:220*, VAL A:324*, GLU A:282 |
| C_47 | -3 | THR A:283, CYS A:275, ASN A:221, VAL A:332, LEU A:331*, MET A:220*, LEU A:321*, VAL A:324*, MET A:320, THR A:279 |
| C_57 | -1 | CYS A:276*, LEU A:321*, ILE A:317*, VAL A:324*, LEU A:331*, MET A:220*, TYR A:334 |
| C_58 | 5.2 | THR A:283, CYS A:275, ALA A:333, CYS A:276*, ILE A:317*, VAL A:332, LEU A:321*, VAL A:324*, MET A:330, MET A:320, MET A:220* |

Interactions common with the control\*

High gastrointestinal absorption was predicted for 31 of the 76 compounds, 13 were predicted to have blood-brain barrier permeant capabilities, and 19 were predicted to be P-glycoprotein substrates. 18 drugs in total were predicted to be CYP-inhibitors of various sorts, of which 6 were predicted to inhibit CYP1A2, 2 CYP2C19, 11 CYP2C9, 5 CYP2D6, and 9 CYP3A4. These results are illustrated in **Table 6**.

**Table 6:**
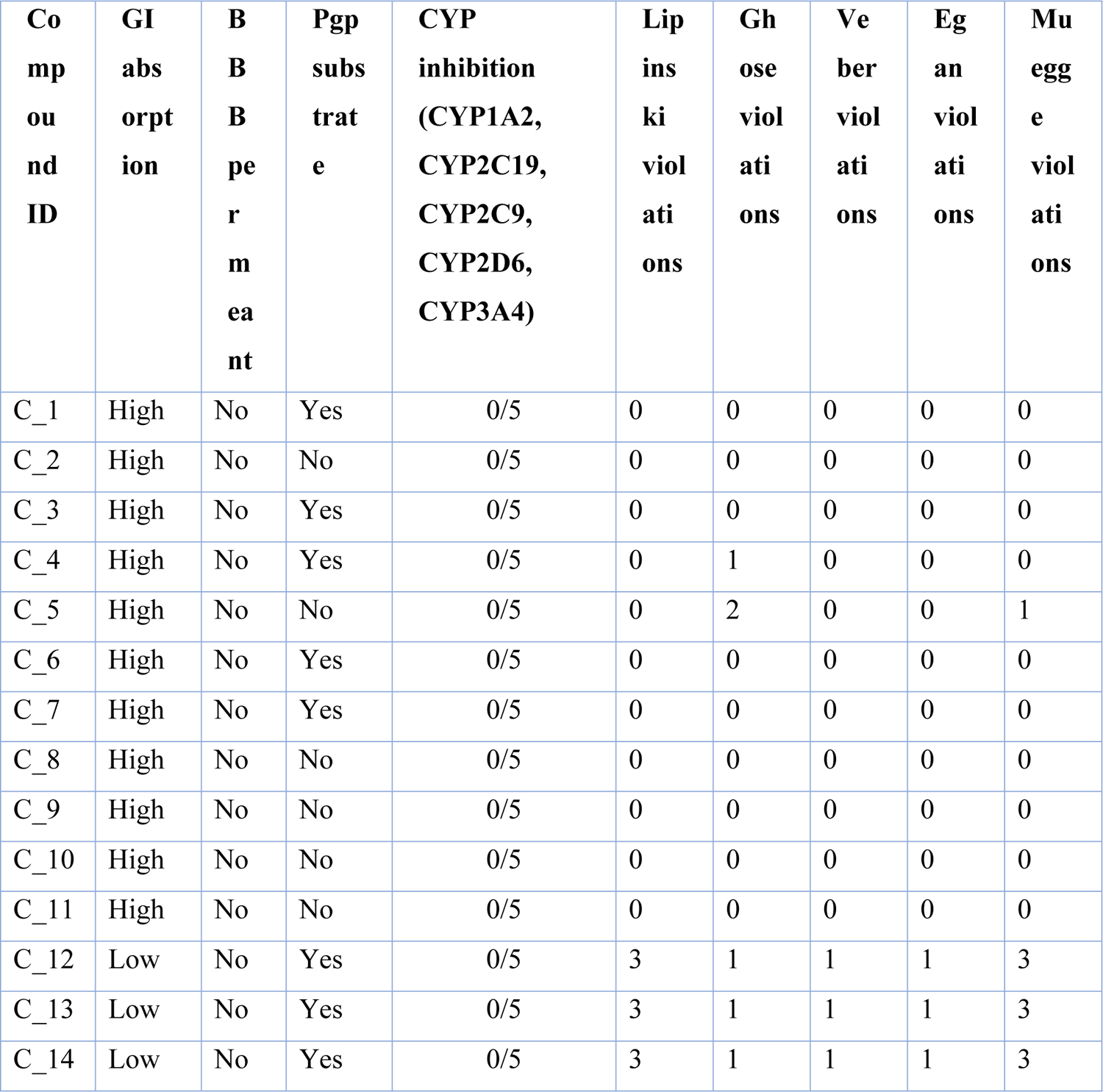
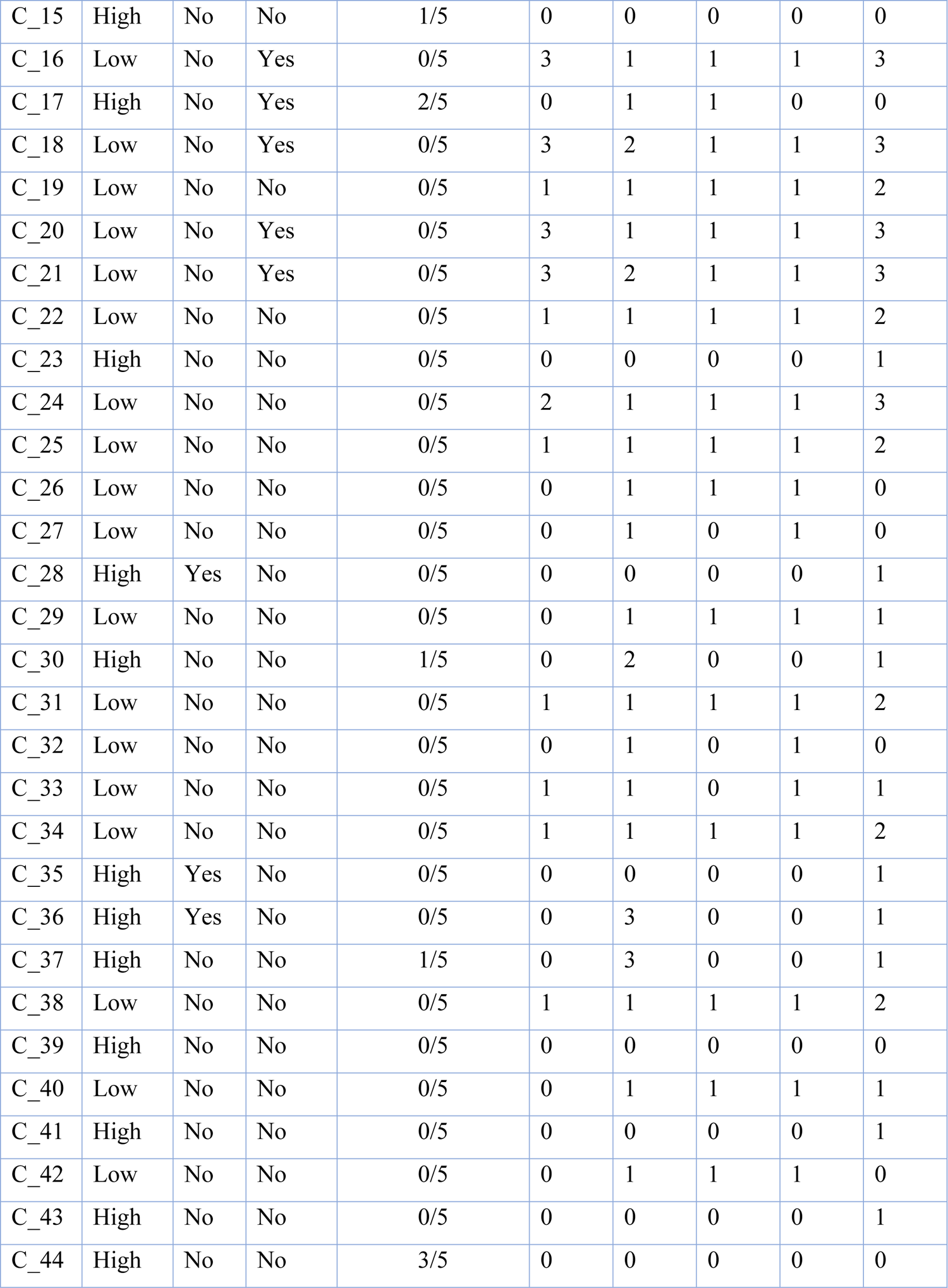
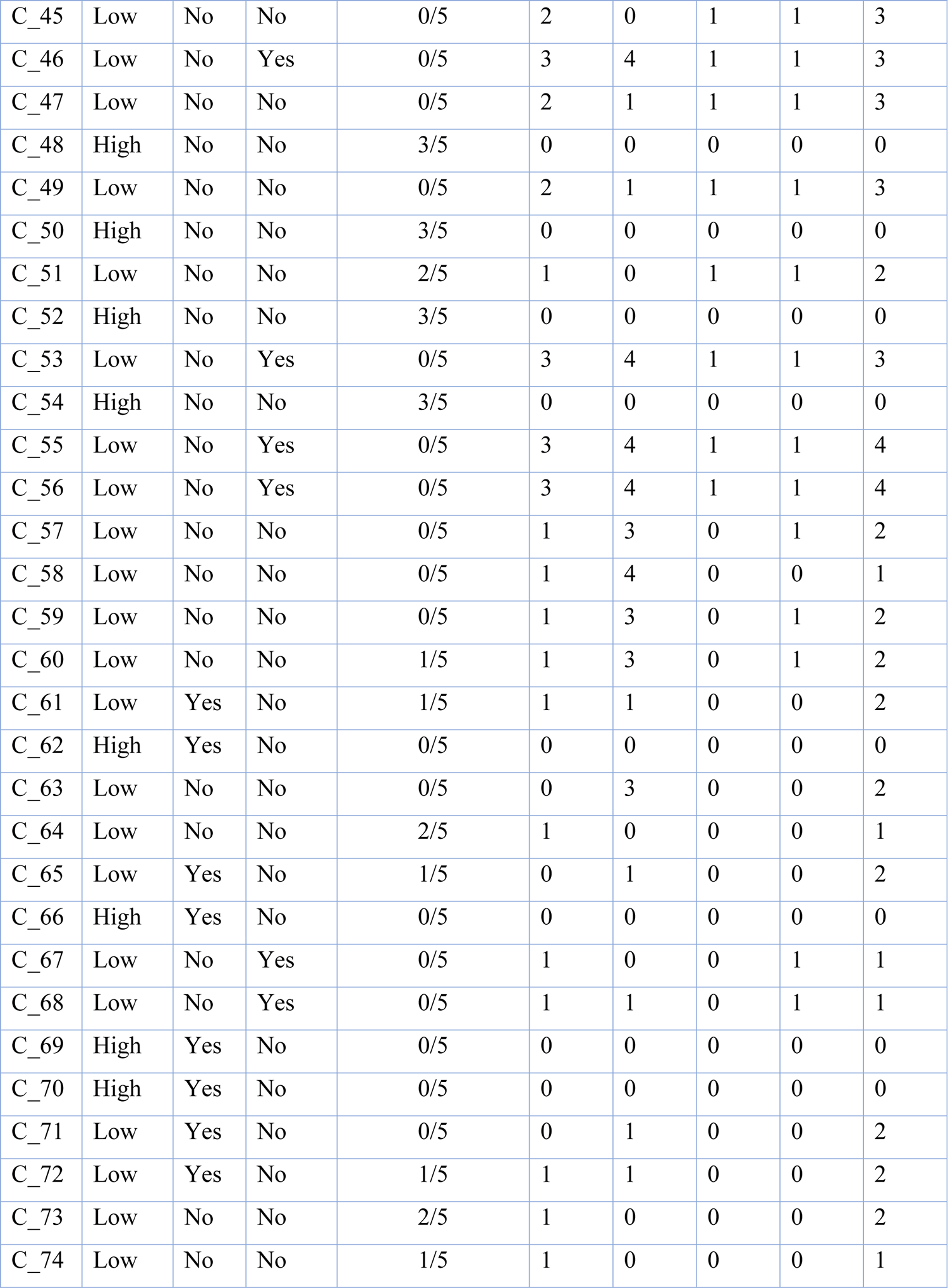
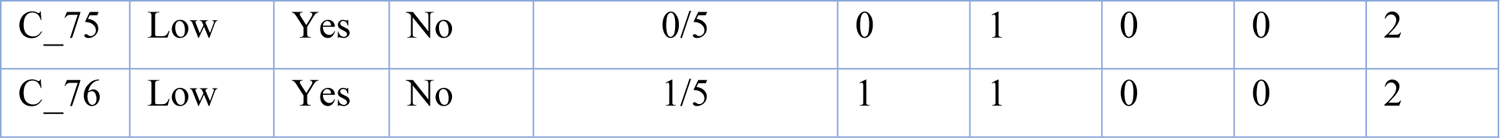
ADME properties of the phytoconstituents

Of the 76 phytoconstituents, only 20 were predicted not to induce any sort of toxicity. 12 were predicted to be highly carcinogenic, 36 highly immunotoxic, and 2 highly mutagenic. Moderate probability of carcinogenicity, mutagenicity, immunotoxicity, and cytotoxicity was observed in 10, 6, 2, and 1 compound(s) respectively. 5 compounds belonged to toxicity class II (fatal if swallowed), 8 to class III (toxic if swallowed), 13 to class IV (harmful if swallowed), and their rest to classes V and VI. These data are presented in **Table 7**.

**Table 7:**
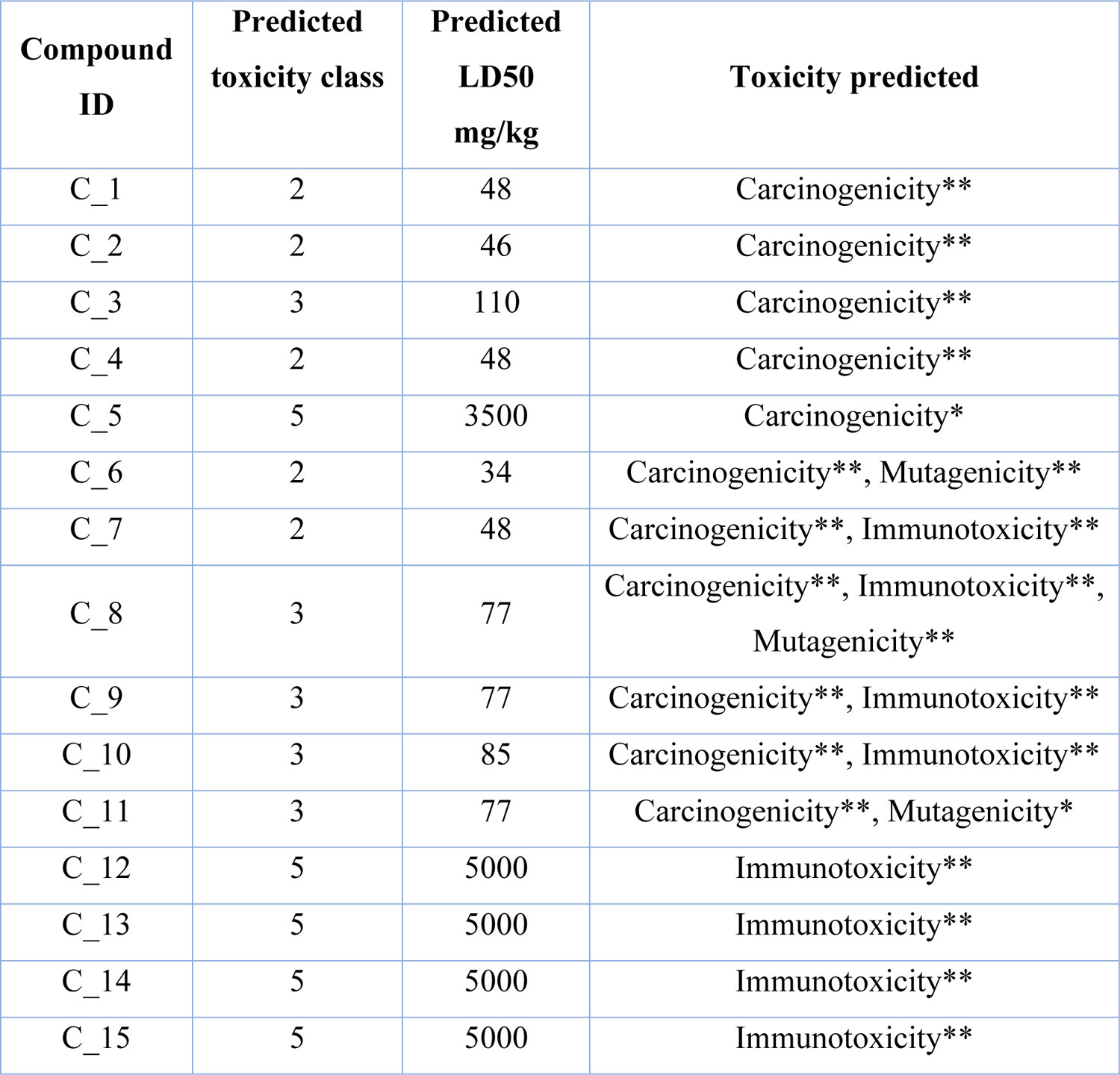
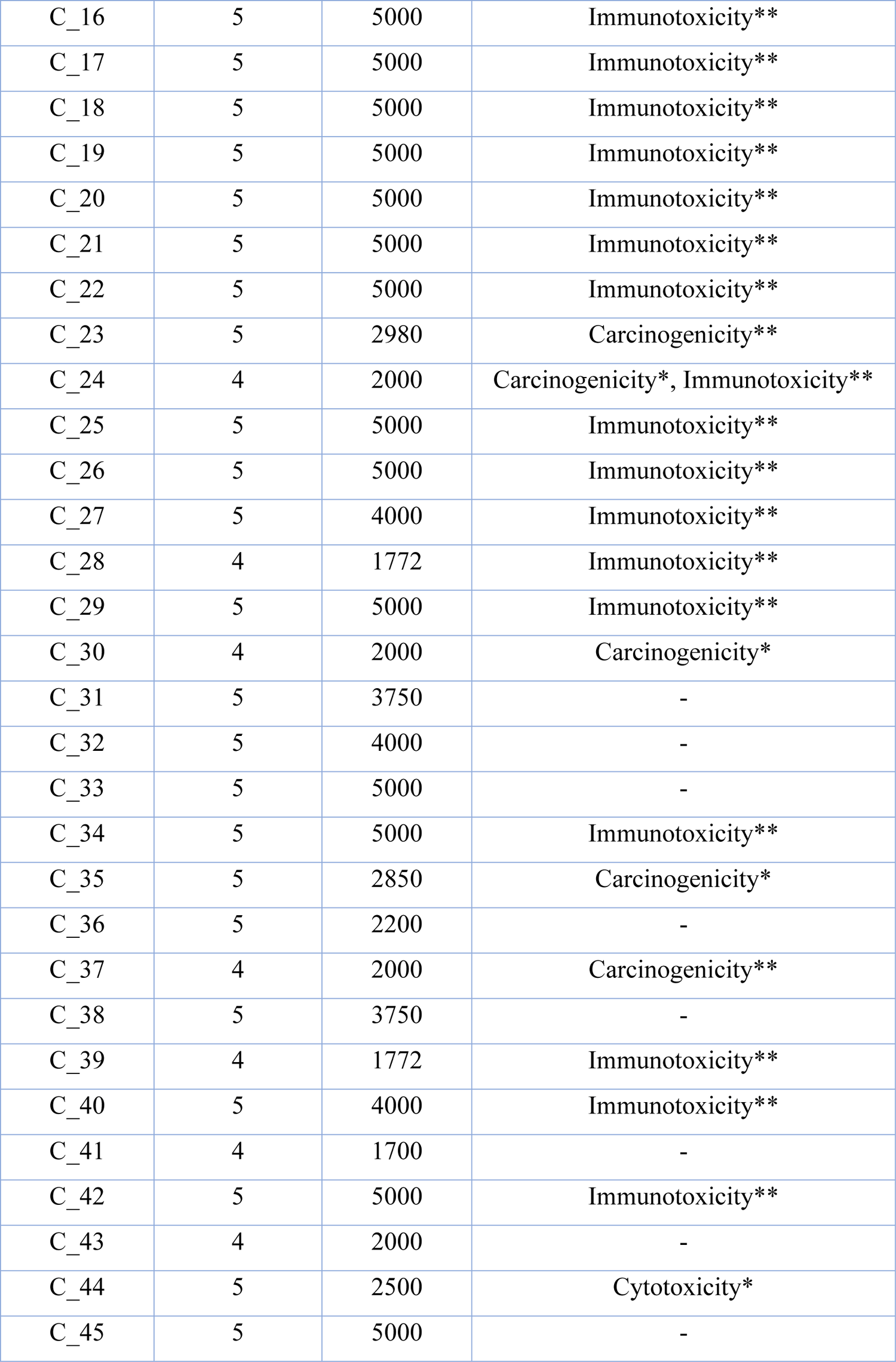
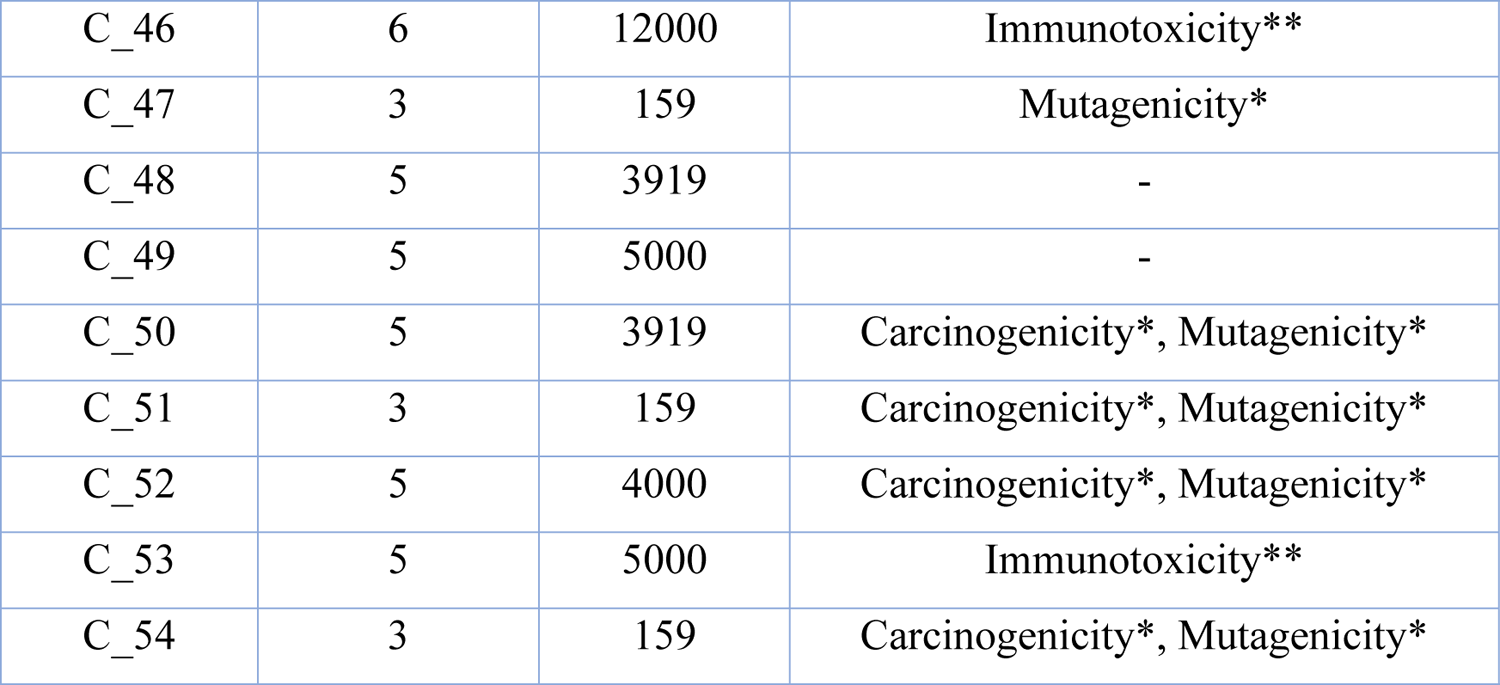
Toxicity profile of the phytoconstituents (Toxicity Class I: fatal if swallowed, Class II: fatal if swallowed, Class III: toxic if swallowed, Class IV: harmful if swallowed, Class V: may be harmful if swallowed, Class VI: non-toxic)

## Discussion

Hepatotoxicity is a critical public health concern on a global scale. As the body’s most preeminent organ, the liver performs a wide range of essential tasks, including secretion, storage, metabolism and detoxification of potentially harmful chemical substances [52–53]. Therefore, any form of hepatic impairment triggered by hepatotoxic agents may carry grave ramifications, culminating in hepatic failure and death. Medicinal plants are a goldmine of potentially therapeutic novel natural substances that can be a puissant sword for combating several liver disorders [54]. The present study investigated the magical hepatoprotective potentialities of *G. procumbens* plant extract against the toxic chemical CCl_4_ that induced hepatotoxicity in Wistar albino male rats via generation of highly reactive free radicals.

The experimental data of the current study demonstrated a drastic reduction in the body weight of rats belonging to the disease control group following exposure to carbon tetrachloride, indicating the occurrence of carbon tetrachloride induced hepatic impairment that may impede the body’s ability to process nutrients, leading to metabolic imbalance and severe loss of body weight. On the other hand, rats in the negative control group showed an increase in terminal body weight, indicating a healthy metabolic homeostasis that regulated the normal growth rate and body weight of rats. However, groups solely treated with the test extract exhibited non-significant (p>0.05) statistical difference in the body weight of rats when compared to the negative control group, inferring that the extract had no detrimental effect on the growth rate of healthy rats; whereas only silymarin-treated groups demonstrated the opposite result, indicating its detrimental effect on the growth rate and body weight of healthy rats. Treatment with 50% ethanolic extract of *G. procumbens* significantly (p<0.05) reversed the CCl_4_-induced loss of body weight in a dose-dependent manner, evincing the promising potency of the extract in altering metabolic imbalance and thus restoring normal growth rate and body weight of diseased rats. Similar findings were reported by Algariri et al. (2013) on the recovery of body weight in diseased rats upon treatment with 50% ethanolic extract of *G. procumbens* [55]. Our findings also showed similarity with those of Islam et al. (2019), who noticed the efficacy of *G. procumbens* ethanolic extract in preventing body weight loss in diseased rats [56]. According to another study conducted by Tahsin et al. (2022), therapy with *G. procumbens* ethanolic extract resulted in pretty consistent effects in recovering body weights of diseased rats, despite the difference that the study was performed using 70% ethanolic extract [57]. Additionally, our study results were in line with those of Ziaul Amin et al. (2021), who found that both the aqueous and 90% ethanolic extracts of *G. procumbens* were significantly effective in regaining body weights in diseased rats in a dose-dependent manner [58].

In figure 2, 3 and 4, a noticeable discrepancy was detected between the two control groups (i.e., negative control group and disease control group) in terms of the levels of some serum enzymes (such as SGPT, SGOT and ALP) that serve as the most sensitive biomarkers of hepatic impairment. In the negative control group, the serum levels of these enzymes were reported to be normal which denoted that the bulk of these enzymes were confined to the liver cells, not in the bloodstream and that was a stark sign of intact, healthy and well-functioning liver cells. On the other hand, a significantly pronounced escalation was noted in these serum enzyme levels of the disease control group, indicating the pathological manifestation of CCl_4_-induced hepatotoxicity, due to which the enzymes (SGPT, SGOT and ALP) leaked out of the liver cells and released into the bloodstream, resulting in elevated serum enzyme levels. However, in-vivo administration of *G. procumbens* to the disease control group was found to cause a dose-dependent dramatic reversal of the CCl_4_-induced alterations in the levels of the hepatic damage biomarkers (i.e., SGPT, SGOT, ALP), yielding statistically non-significant (p>0.05) outcomes with the standard drug (silymarin). This evinced the promising hepatoprotective action of *G. procumbens*. The serum enzyme levels of the remaining six groups (non-CCl_4_ treated) demonstrated statistically non-significant (p>0.05) deviation from those of the negative control group, signifying the potential safety margin of silymarin and *G. procumbens*. Our findings regarding the serum SGPT and SGOT reversal efficacy of *G. procumbens* ethanolic extract were quite consistent to those of Islam et al. (2019) and Tahsin et al. (2022), who found significant serum enzyme (i.e., SGPT and SGOT) restoration efficacy of G. procumbens employing 100% and 70% ethanolic extract respectively [56–57].

Creatinine is considered to be a fairly accurate indicator of renal function. Figure 5 depicted a notable increment in creatinine levels in diseased rats, denoting reduced renal efficacy in clearing creatinine from the bloodstream. This may be due to hepatotoxicity associated physiological alterations that adversely affected the renal perfusion and blood circulation, resulting in impaired kidney function. But the creatinine level was reported to be normal in non-diseased individuals (negative control group), signifying healthy kidney functioning. The values of creatinine levels recorded from the diseased rats treated with three different doses (low, medium and high) of *G. procumbens* were significantly lower than the values obtained from the disease control group, illustrating the dose-dependent efficacy of *G. procumbens* in attenuating the risk of hepatotoxicity-induced kidney damage. Though this study was performed with 50% ethanolic extract of G. procumbens, our results are strikingly comparable to those of Tahsin et al (2022) and Ishak et al (2021), both of whom used a 70% ethanolic extract of the plant, reporting effects that were quite identical to ours in terms of restoring creatinine levels in diseased rats [58, 59]. Our observations also showed consistency with those of Islam et al. (2019), who noticed a significant efficacy of *G. procumbens* in balancing serum creatinine levels of diseased rats with 100% ethanolic extract of the plant [56].

As illustrated in figure 6, the serum LDH level of the negative control group was strikingly inconsistent with that of the disease control group, where the serum LDH level was archly elevated upon treatment with CCl_4_ that indicated the occurrence of enzyme leakage into the bloodstream caused by hepatotoxicity-induced demolition of hepatocytes. But the serum LDH levels of the six non-CCl_4_ treated groups showed non-significant (p>0.05) statistical deviation from those of the negative control, portending the absence of any hazardous risks or fatal effects associated with the consumption of silymarin or *G. procumbens*. However, a dose-dependent curtailment in serum LDH level was noted after treating the diseased rats with differing doses of *G. procumbens*, evincing the pronounced action of *G. procumbens* in normalizing the serum LDH level by repairing the damaged hepatocytes caused by CCl_4_-mediated hepatotoxicity. These results were fairly similar to those of Kim et al (2006), Halim et al (2021) and Tahsin et al (2022) who reported the efficacy of *G. procumbens* in normalizing serum LDH level employing aqueous extract, 80% ethanolic extract and 70% ethanolic extract respectively [57], [60–61].

According to figure 7, 8, 9 and 10, the levels of serum lipids (i.e., HDL, LDL, triglyceride and total cholesterol) of the negative control group were within the normal range, denoting homeostasis in serum lipid profiles of healthy rats. But in-vivo induction of CCl_4_ significantly affected the serum lipid profiles of the treated rats, resulting in a considerable increase in the levels of serum LDL, total cholesterol and triglyceride with a substantial drop in serum HDL level. Such alterations in serum lipid levels of the CCl_4_-treated rats might be attributable to a malfunctioning hepatic de novo lipogenesis pathway, hinting at the occurrence and pathological progression of CCl_4_-prompted hepatic impairment. But treatment with *G. procumbens* notably reversed the CCl_4_-induced alterations in serum lipid profile in a dose dependent manner, providing non-significant (p>0.05) statistical outcomes with the standard medication (silymarin), which proved the potential antihepatotoxic action of *G. procumbens*. Groups solely treated with the standard (silymarin) or test extract *(G. procumbens)* imitated the same trend as the negative control group, evincing the impossibilities of potentially harmful side effects following intake of silymarin or *G. procumbens*. Consistent with our observations, Tahsin et al. (2022) and Ahmad Nazri et al. (2019) reported a significant serum lipid profile-balancing efficacy of *G. procumbens* in diseased rats, in spite of the contrariety that they performed the experiments utilizing 70% and 80% ethanolic extract of the plant respectively [57, 62]. Compared to our investigations, another study conducted by Ziaul Amin et al. (2021) reported that both of the aqueous and 90% ethanolic extracts of *G. procumbens* had impressive effects on restoring the altered levels of serum cholesterol, triglyceride and HDL in diseased rats, but the aqueous extract exhibited slightly superior outcomes compared to the ethanolic one [58]. However, in contrast to our findings, they detected no significant alterations in LDL levels of the diseased rats upon treatment with differing doses of aqueous and ethanolic extracts of *G. procumbens* [58]. This may be owing to the fact that the plant’s particular constituent, which helped restore the LDL level, may not have a favorable environment to grow in sufficient quantities due to differences in geographical region or soil composition. Another possible explanation may be that their study was performed with different solvent compositions than ours, which may have contributed to the reduction of extraction efficacy of *G. procumbens*, resulting in a loss of bioactive elements that could aid in the restoration of LDL levels in diseased rats.

In figure 11, there was no evidence of DNA fragmentation in the negative control group, implying that the DNA was completely intact; while in the disease control group, a significantly evident increase in DNA fragmentation was detected, signifying an early indication of apoptosis. However, the plant extract was found to significantly attenuate the CCl_4_-induced elevation in DNA fragmentation in the disease control group in a dose dependent manner, denoting its efficacy in mending apoptosis. These results yielded by the test extract were statistically non-significant (p>0.05) when compared to those of the standard drug, silymarin. The DNA fragmentation profiles of the non-CCl_4_ treated groups were pretty consistent with that of the negative control group, inferring that neither silymarin nor *G. procumbens* ingestion would cause any discernible deleterious effects on the DNA fragmentation profiles of the healthy individuals.

**Figure 11:**
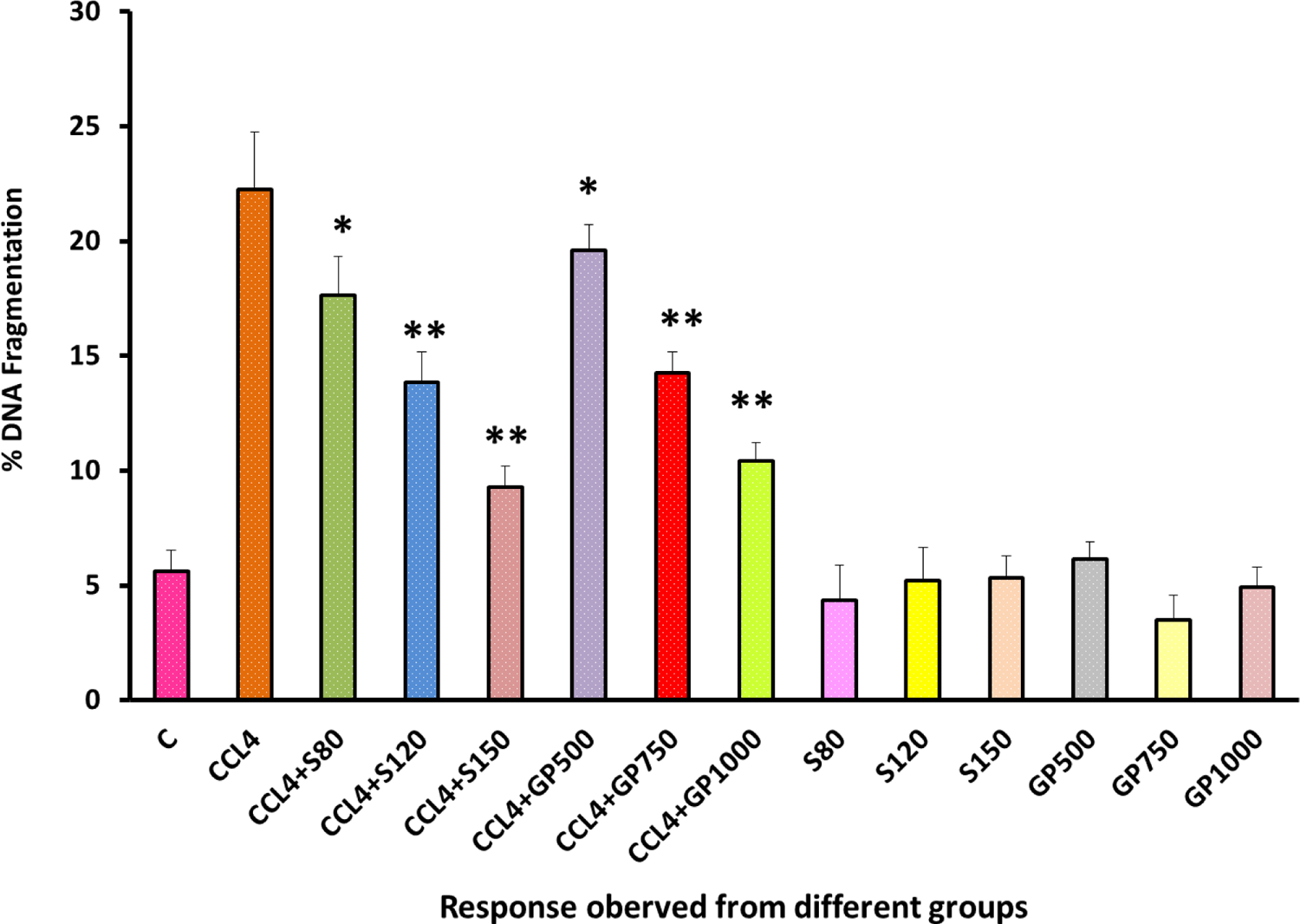
DNA fragmentation Level of rats from 14 groups. The data were expressed as mean ± standard deviation. (* indicates statistically significant change)

According to figure 12, the serum level of gamma glutamyl transpeptidase (γGT) enzyme was found to be normal in the negative control group, which implied that the majority of the enzyme was stored within the liver cells, indicating healthy liver functioning. But in the disease control group, the serum level of this enzyme was significantly (p<0.05) elevated. This may be a result of CCl_4_-induced hepatic impairment, which may have caused the γGT enzyme to seep out of the hepatocytes and be released into the bloodstream, leading to a rise in serum γGT levels. But treating the diseased rats with differing doses (e.g., low, medium and high) of silymarin or *G. procumbens* significantly lowered the elevated levels of serum γGT enzyme. Though silymarin-treated groups exerted this effect at a slightly higher magnitude compared to the extract-treated groups, both of them generated statistically non-significant (p>0.05) results with the negative control group at their highest doses which evinced the dose-dependent efficacy of silymarin and *G. procumbens* in restoring the altered level of serum γGT enzyme. The serum γGT level of the six non-CCl_4_ treated groups demonstrated statistically non-significant (p>0.05) deviation from those of the negative control group, signifying the absence of potential menace upon intake of silymarin or *G. procumbens*. Our findings regarding the serum γGT level restoration efficiency of *G. procumbens* were quite similar to those of Ismail et al. (2016), despite the difference that they performed their experiment employing aqueous extract of G. procumbens [63].

**Figure 12:**
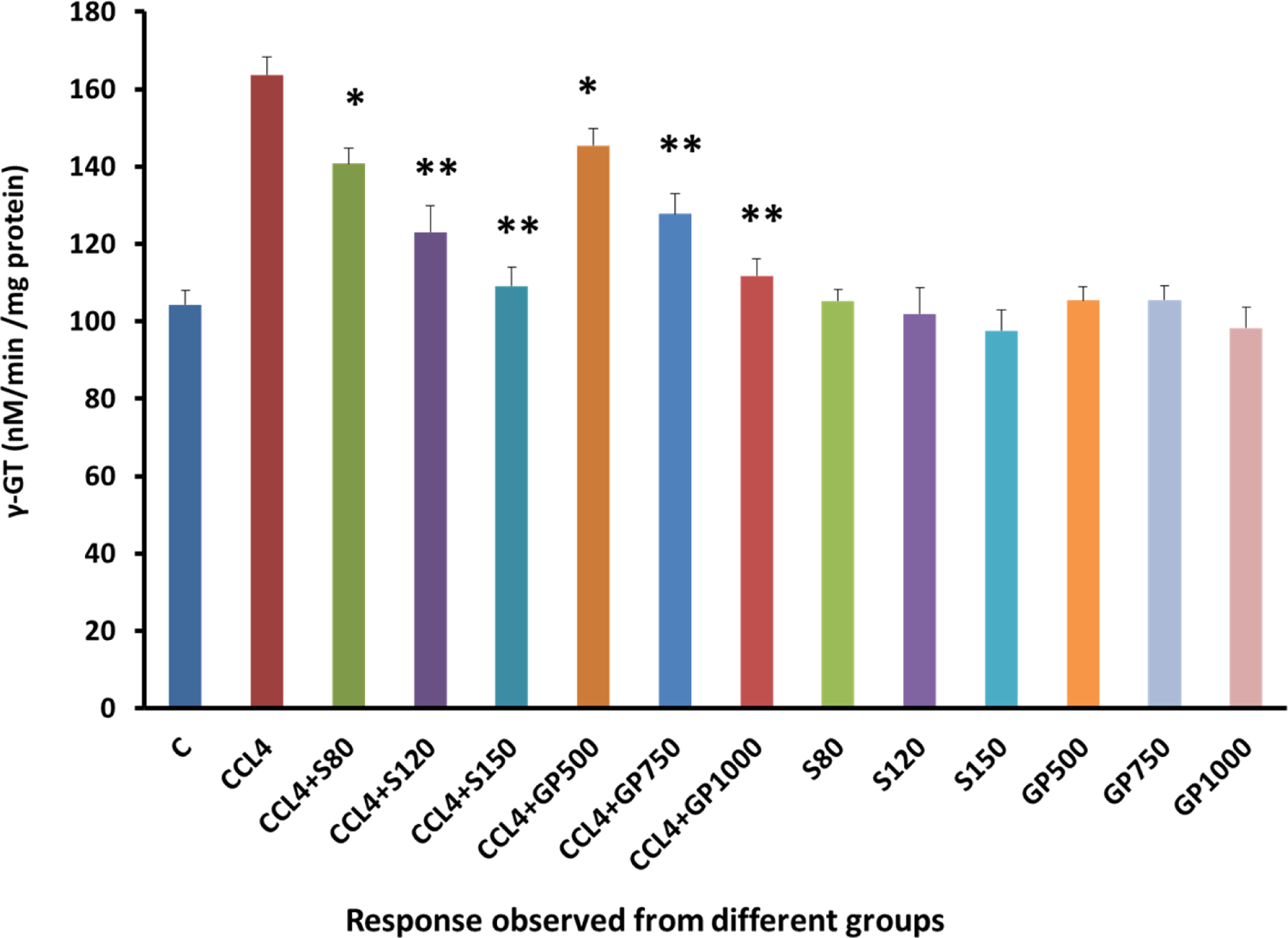
γGT (nM/min/mg protein) Level of rats from 14 groups. The data were expressed as mean ± standard deviation. (* indicates statistically significant change).

**Figure 13:**
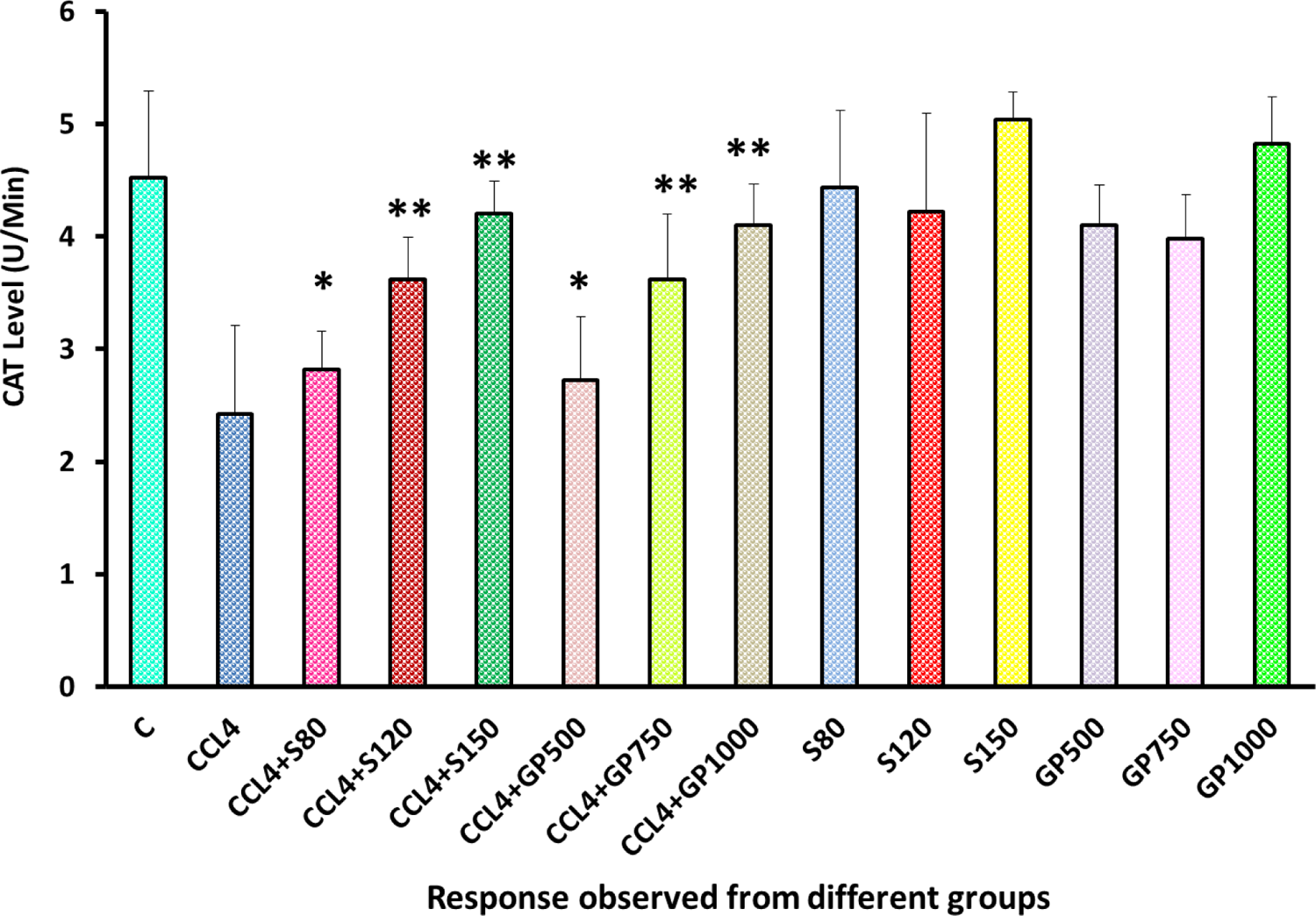
CAT (U/Min) Level of rats from 14 groups. The data were expressed as mean ± standard deviation. (* indicates statistically significant change)

**Figure 14:**
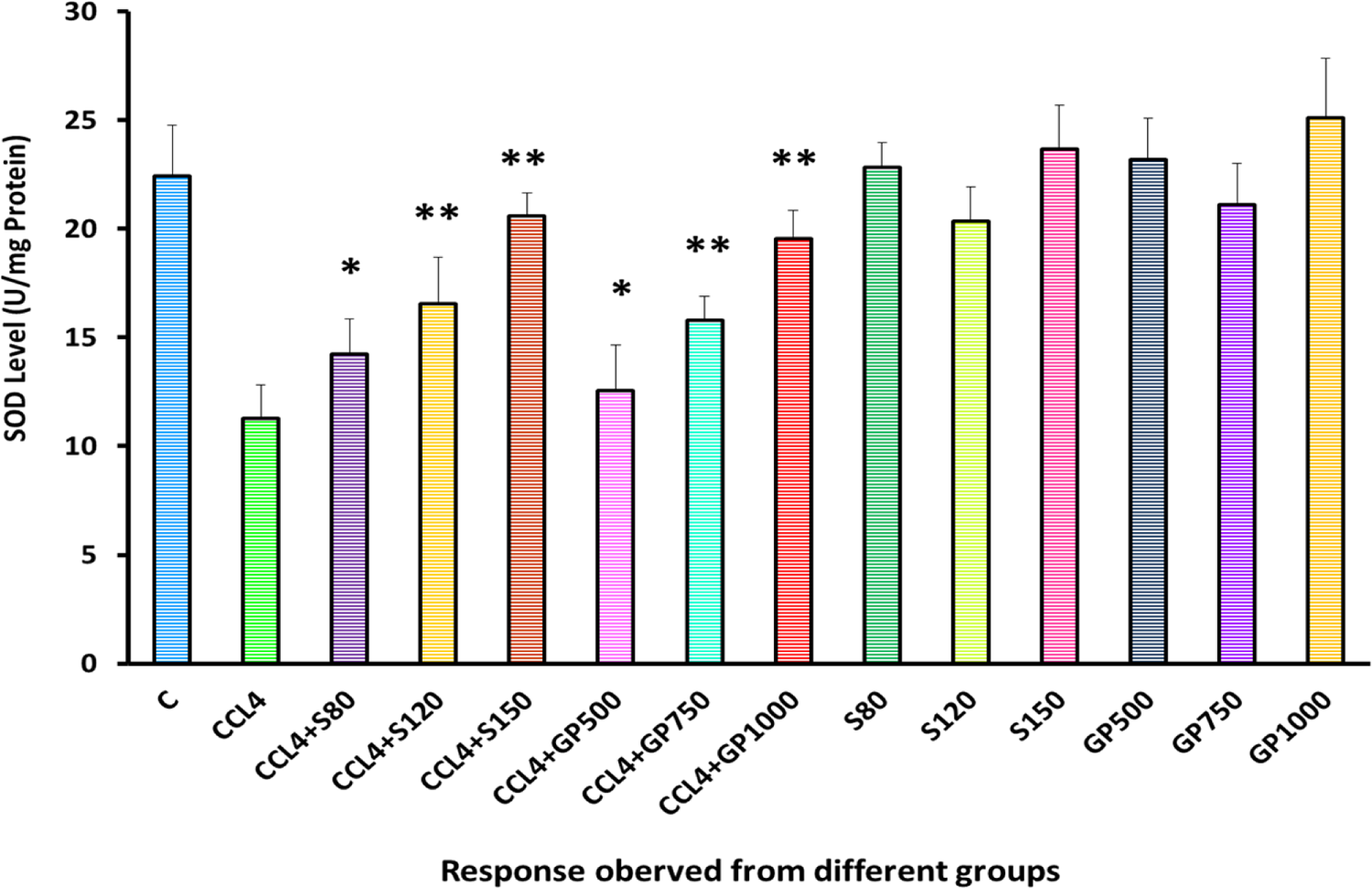
SOD (U/mg Protein) Level of rats from 14 groups. The data were expressed as mean ± standard deviation. (* indicates statistically significant change)

**Figure 15:**
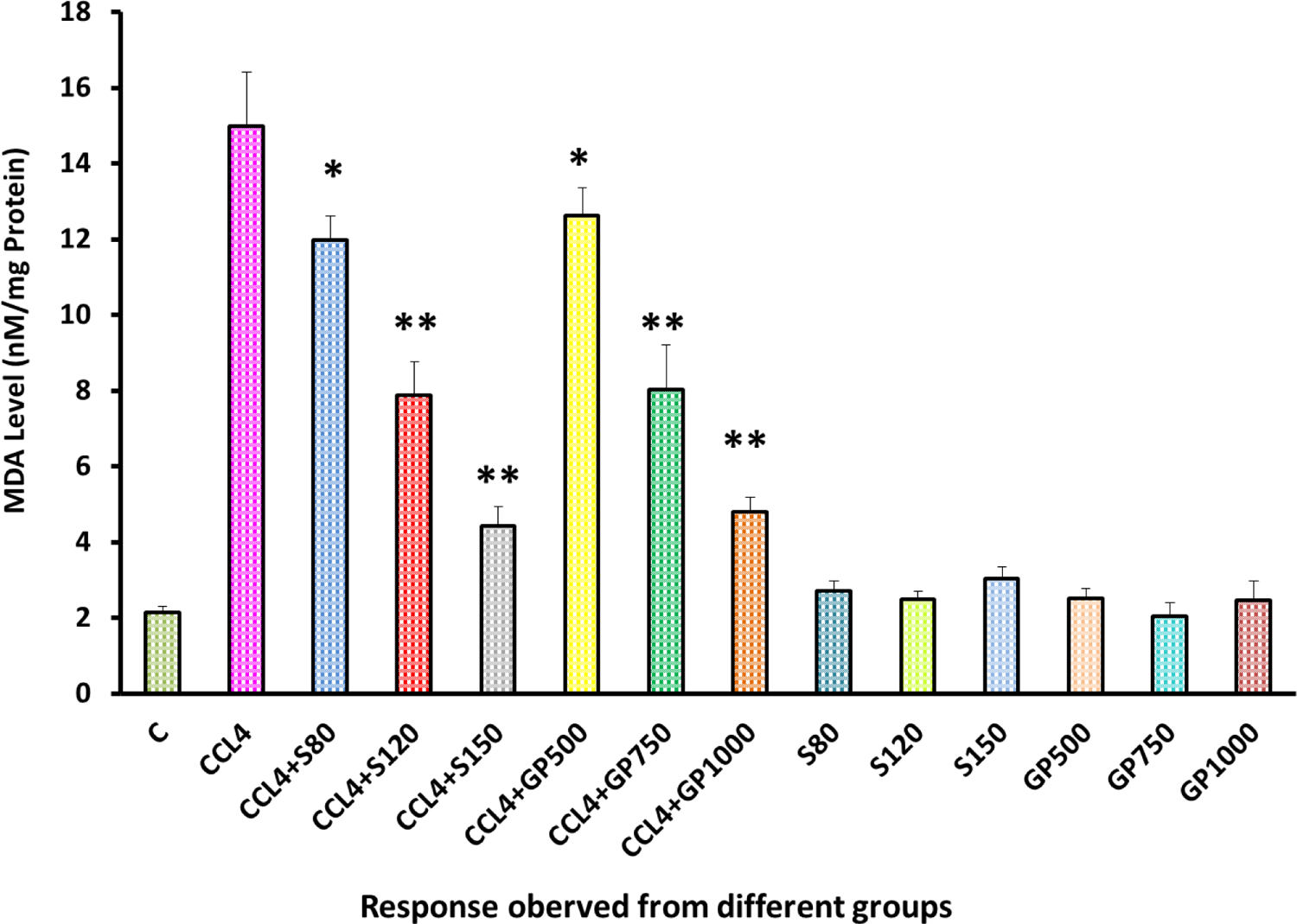
MDA (nM/mg Protein) Level of rats from 14 groups. The data were expressed as mean ± standard deviation. (* indicates statistically significant change)

**Figure 16:**
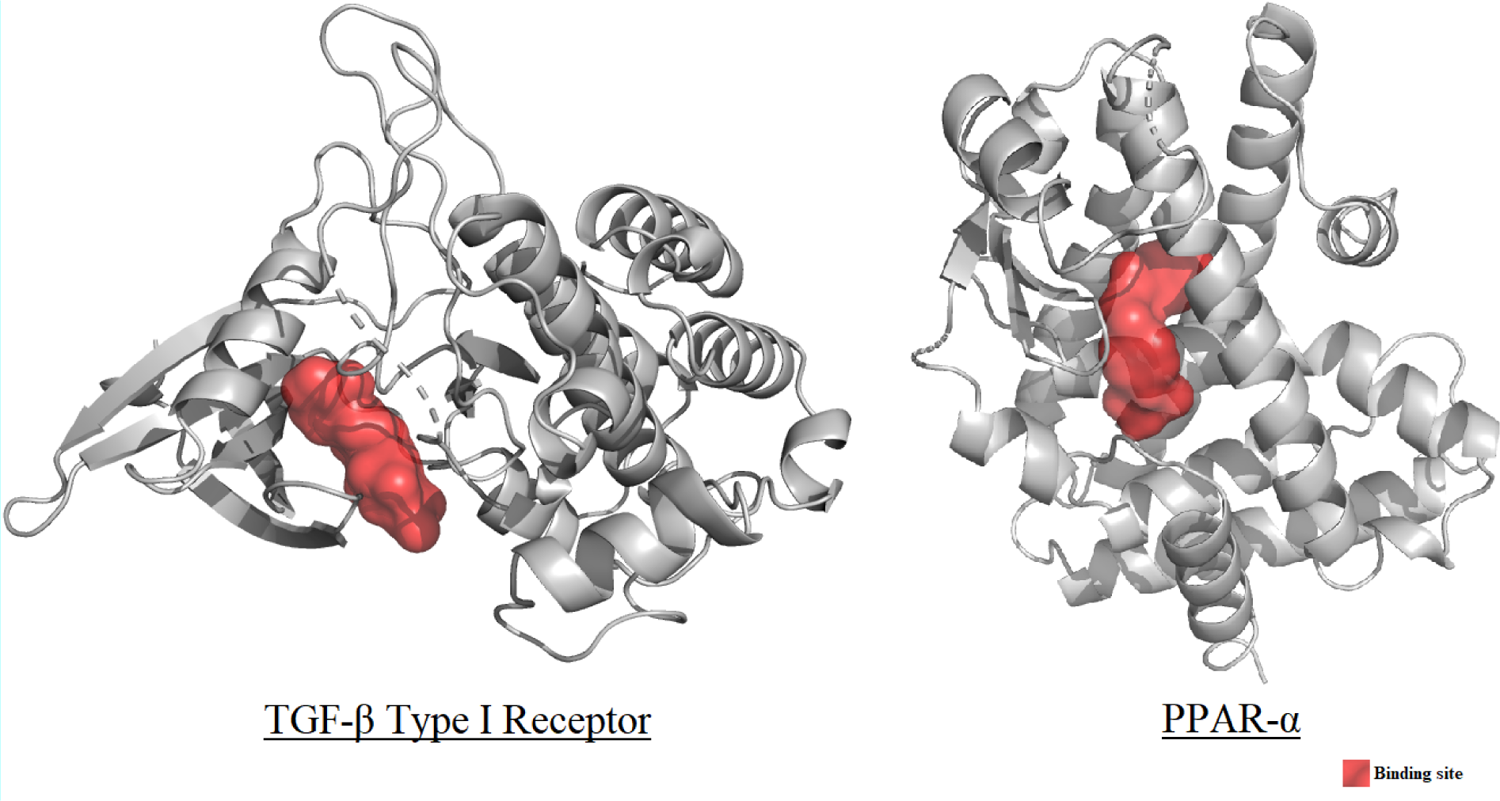
Binding sites of macromolecules TGF-β type I receptor and PPAR-α (binding site marked in red)

**Figure 17:**
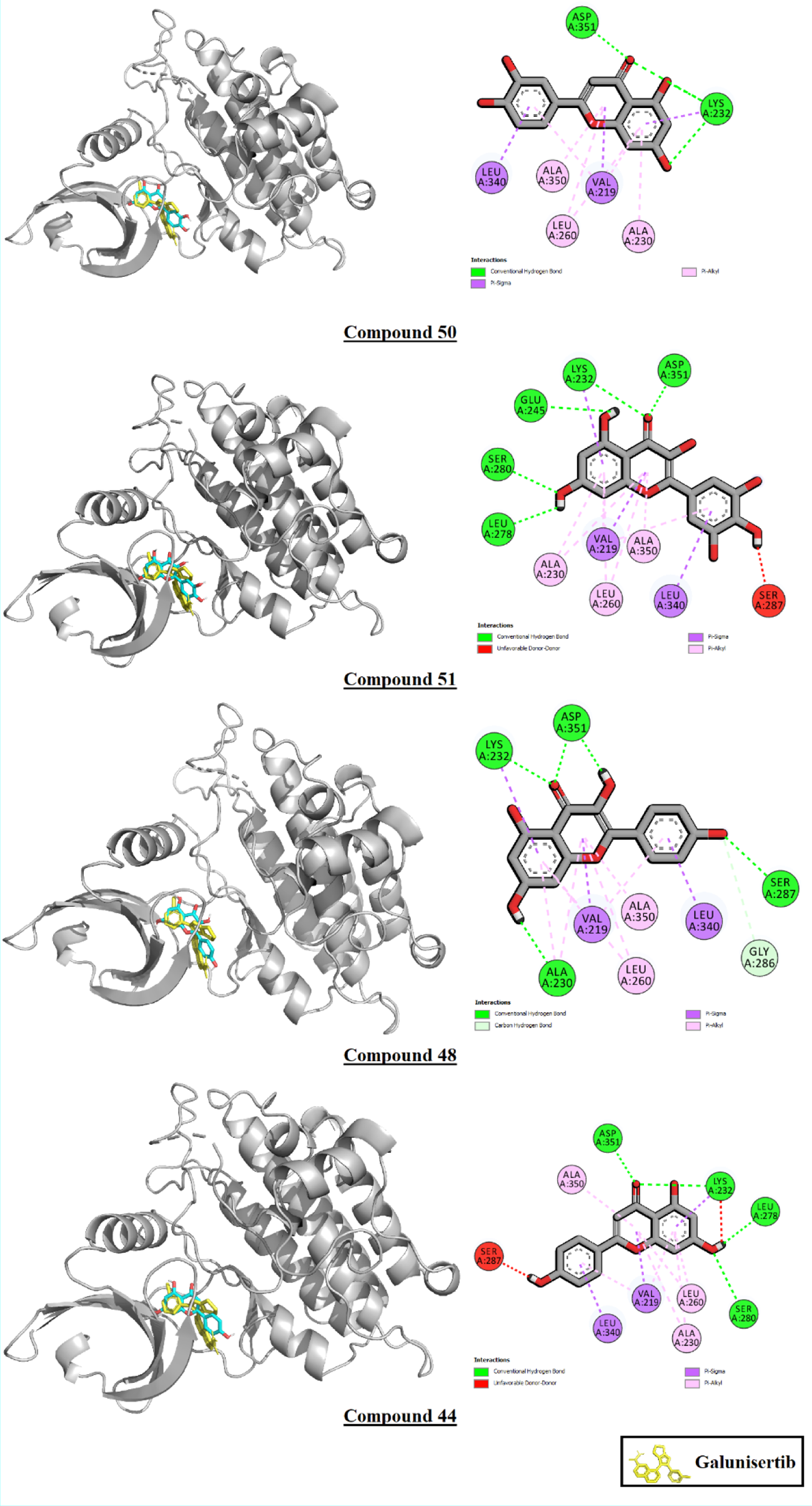
Interactions of compounds 44, 48, 50, and 51 with TGF-β type I receptor

**Figure 18:**
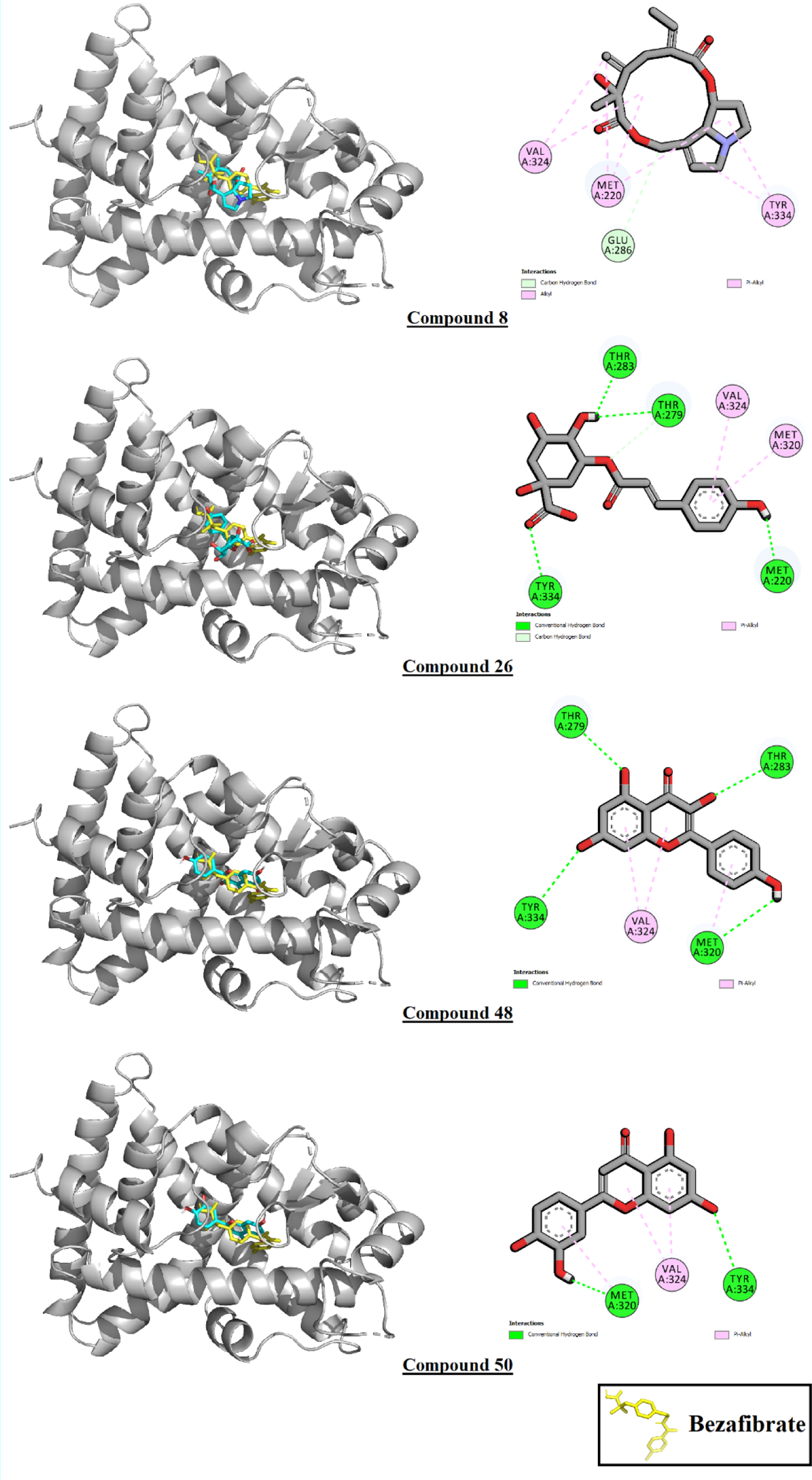
Interactions of compounds 8, 26, 48, and 50 with PPAR-α

Hepatic impairment was found to be intricately associated with oxidative stress [64]. CAT, SOD and MDA serve as biomarkers of oxidative stress. Upon treatment with carbon tetra chloride, a significant (p<0.05) diminution was observed in the levels of SOD (superoxide dismutase) & CAT (catalase) enzymes in conjunction with a significant increase (p<0.05) in MDA (malondialdehyde) levels of diseased rats, illustrating a sharp disparity with those of the healthy individuals. The alterations in the levels of these enzymes could be a result of the enhanced production of free radicals by the toxic substance, CCl_4_. The presence of low serum levels of SOD and CAT enzymes supported the premise that the defensive actions of these antioxidants were diminished, resulting in increased oxidative damage. On the other hand, elevated levels of serum MDA denoted higher oxidative stress, more specifically higher lipid peroxidation that may act as a contributing factor in the pathophysiology of hepatic impairment. However, administration of fluctuating doses (low, medium and high) of *G. procumbens* to the disease control group was noticed to cause an effective reversal in CCl_4_-induced alterations in the levels of these enzymes (SOD, CAT, MDA), generating statistically non-significant (p>0.05) outcomes with the standard drug silymarin that evinced the dose dependent efficacy of *G. procumbens* in attenuating CCl_4_-induced oxidative stress. Though our study was performed with 50% ethanolic extract of *G. procumbens*, our findings are noticeably comparable to those of Halim et al. (2021) and Ahmad Nazri et al. (2019), both of whom conducted their studies utilizing 80% ethanolic extract of the plant, reporting effects that were pretty consistent to ours in terms of restoring the levels of oxidative stress biomarkers (CAT, SOD and MDA) in diseased rats [61–62].

Furthermore, the findings of the current study revealed the superiority of *G. procumbens* in minimizing some of the side effects of antihepatotoxic drugs that are currently available on the market. Many commercially available drugs used for the treatment of various hepatic disorders showed hepatotoxic effect in several cases. For instance, Lactulose-a nonabsorbable disaccharide is considered as the standard treatment option for hepatic encephalopathy [65]. Dietary lactulose (5%, w/w) is known to increase total cholesterol levels in female rats of an outbred Wistar strain by influencing cholesterol metabolism [66]. In another study, the effects of lactulose on the liver in treatment of obstructive jaundice was investigated in rat model and serum liver enzymes (AST, ALT, γ-GT, and ALP) & MDA levels were found to be significantly higher [67]. In our study, *Gynura procumbens* leaf extract significantly decreased serum total cholesterol, γ-GT, ALP & MDA levels in rats when administered orally.

Drug induced liver injury (DILI) is considered one of the common causes of withdrawal of approved medications from the marketplace. Sofosbuvir as monotherapy or in combination with Ribavirin is indicated for the treatment of chronic hepatitis C infection [68]. Among all hospitalized DILI cases in a tertiary Egyptian center, Sofosbuvir plus ribavirin dual therapy was reported to cause 2.7% drug induced liver injury with elevated ALT more than 3-fold and/or ALP more than 2-fold the upper limit of normal value [69]. In contrast, our current findings indicate that 50% ethanol extract of Gynura procumbens leaves reduced serum ALP levels better than negative control group and ensured the effectiveness of the extract alone in treating CCl4 induced hepatotoxicity in rats.

Furthermore, in small doses some drugs may act as antioxidants whereas sometimes may act as pro-oxidants in high doses. According to Ibrahim et al. (2018) high dose (82.2 mg/kg) of Sofosbuvir did not significantly improve thioacetamide-induced liver injury in rats [70]. In other words, there was no significant effect on liver enzymes (i.e., ALT, AST, ALP etc.) or MDA content compared with disease control group. Such hepatotoxic effect of Sofosbuvir was attributed to the accumulation of a high concentration of its metabolites. However, results of present study showed statistically significant reduction in serum ALP & MDA levels, even after the administration of extract at higher dose (1000mg/kg).

Long term use of hepatoprotective medication may result in some hepatotoxic effect. Widely used nucleoside analogue antiretroviral drug, Lamivudine is first line of treatment for chronic hepatitis B and also known for its low toxicity at clinically prescribed dose [71]. However, hepatotoxic potentials of lamivudine at high doses (≥ 100 mg/kg) on prolonged administration was observed when hepatic γ-GT and MDA concentration significantly elevated at 20 mg/kg in rats [72]. This finding was inconsistent with our in vivo findings as γGT & MDA levels returned to the same level of healthy group in a dose dependent manner, precisely with last highest dose of *Gynura procumbens* extracts. Another Human model study of long-term combination therapy using ursodeoxycholic acid and bezafibrate in the treatment of primary biliary cirrhosis and dyslipidemia showed significantly increased the serum creatinine levels (mean 0.94 mg/dl). The serum creatinine level gradually increased shortly after the initiation of bezafibrate administration [73]. On the contrary, oral administration of *Gynura procumbens* extracts in rats noticeably decreased serum creatinine level in this study.

Paracetamol hepatotoxicity, with is the predominant cause of acute liver failure in the western countries. McGill et al. (2012) suggested that mitochondrial damage and nuclear DNA fragmentation are likely to be critical events in paracetamol hepatotoxicity in humans, resulting in necrotic cell death [74]. DTS, a novel nutraceutical, showed considerable mitigation of acute DNA fragmentation and increased mitochondrial SOD fraction, thus protected oligonucleosomal DNA integrity in rats with high-dose paracetamol [75]. This observation further corroborates our findings, where DNA fragmentation ranges were significantly dropped and noticeably improved SOD level by plant extracts, thus exerted a significant hepatoprotective potential against CCl_4_ induced liver injury in rats.

For the in-silico assessment of hepatoprotectivity via molecular docking, TGF-β1 and PPAR-α were selected as target macromolecules. TGF-β1 plays a major role in redox imbalance by upregulation of ROS production and the down regulation of the anti-oxidant defense system. TGF-β1 induces growth inhibition and apoptosis in hepatocytes both in vivo and in vitro. Elevated TGF-β1 expression indicates onset of fibrosis/cirrhosis. TGF-β1 is also a strong apoptosis inducer and its signaling is closely related with acute liver injury [33]. According to Dooley et al., blocking TGF-β1 signaling pathway by ectopic expression of Smad7 in hepatocytes efficiently inhibited CCl_4_-dependent liver damage and fibrogenesis in mice. Similarly, Proper function of the peroxisome proliferator-activated receptor alpha (PPARα) is essential for the regulation of hepatic fatty acid metabolism [34]. Dysregulation of this gene or its downstream targets has also been implicated in liver-specific disease states. For instance, PPAR-α downregulation is linked to inhibition of HCV replication [35]. Thus, Inhibition of proteins TGF-β1 and PPAR-α has been proposed as the likely mechanisms for hepatoprotective effect [36].

In the molecular docking analysis, G. procumbens phytoconstituents performed poorly against TGF-β1 compared to the control drug galunisertib. Galunisertib scored a binding affinity value of −11.5 Kcal/mole, while the highest scoring flavonoid, compound 50, or luteolin, scored −10.3. The highest scoring phenolic compound, compound 20, or 4, 5-dicaffeoylquinic acid, scored −10.1. This difference may be owed to the structural disparities between these molecules and their subsequent abilities to occupy the binding pocket of the macromolecule.

Panel (2) Galunisertib, Compound 50 and Compound 20 superimposed in their binding poses with the macromolecule TGF-β1. Galunisertib has been marked in yellow, Compound 50 in pink and Compound 20 in green.

From figure 19 panel 2, the steric disparity between the three molecules is apparent. Galunisertib has two conjoined pentacyclic ring structures, the second of which has two branches attached to it, one of which contains a single hexacyclic ring, and the other two conjoined hexacyclic rings. This unique structure allows galunisertib to form interactions with 12 residues in the binding pocket. While compound 50 has a partially similar steric configuration, it lacks the cyclic side branch, and therefore is unable to occupy the binding pocket to a similar degree as galunisertib. Compound 20’s steric configuration is more akin to that of galunisertib, however, it lacks cyclic structures in a number of positions compared to galunisertib. Moreover, a cyclic protrusion at the junction point contributes little to ligand-macromolecule interactions but likely causes steric hindrance, resulting in a lower binding affinity than the control. Steric similarities are of utmost importance in computational drug design, and increasing the steric similarities through structural modification may result is better binding affinities with TGF-β1 [76–78].

**Figure 19:**
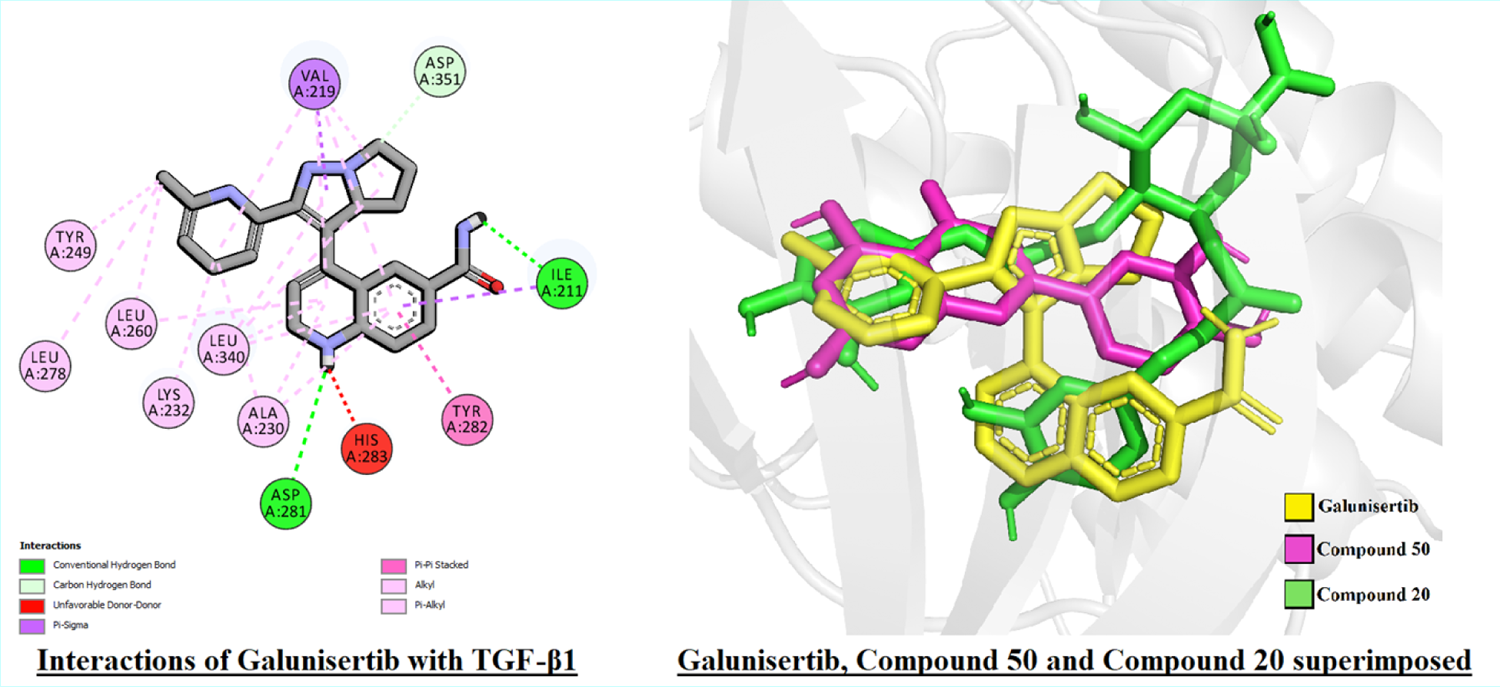
Panel (1) Interactions of Galunisertib with TGF-β1 receptor.

Molecular docking analysis revealed that the phytoconstituents performed considerably better than the control bezafibrate against PPAR-α, with 26 phytoconstituents scoring better than the control (B.A.: −7.6 Kcal/mole). A few high scoring compounds (compounds 25, 26, and 38) did not share any interaction residues with the control, but this was not observed across the library. For both macromolecules, flavonoids and phenolic compounds comprised of a major portion of the top scoring ligands. In case of TGF-β1, all 10 of the best scoring phytoconstituents belonged to either of these two classes. In case of PPAR-α, 9 of the top 10 best scoring phytoconstituents were flavonoids or phenolic compounds. Flavonoids and phenolic compounds have long been known to exert hepatoprotective activity, and this was reflected here as well [79–81]. Overall, the phytoconstituents of *G. procumbens* showed promise as hepatoprotective agents and therefore should be considered strongly for further analysis.

Taken overall, *Gynura procumbens* might be a promising, clinically-applicable, natural counteraction against drug-induced side effects when long term or multiple drug therapy is required for a potent hepatoprotective action.

## Conclusion

Observing the obtained data we can conclude that *G. procumbens* have potential hepatoprotective effects and can be utilized by modifying medicinally to act as a drug against hepatotoxicity.

